# High-resolution magic angle spinning NMR of KcsA in liposomes: the highly mobile C-terminus

**DOI:** 10.1101/2022.06.28.498019

**Authors:** Gary S. Howarth, Ann E. McDermott

**Affiliations:** Department of Chemistry, Columbia University, New York, NY 10031

**Keywords:** solid-state nuclear magnetic resonance (SS NMR), high-resolution nuclear magnetic resonance (HR-MAS NMR), KcsA, liposomes, lipid chemical shifts, lipid hydrolysis, membrane protein structure

## Abstract

The structure of the transmembrane domain of bacterial potassium channel KcsA has been extensively characterized, yet little information is available on the structure of its cytosolic N- and C-termini. This study presents high-resolution magic angle spinning (HR-MAS) and fractional deuteration as tools to study these poorly resolved regions for proteoliposome-embedded KcsA. Using ^1^H-detected HR-MAS NMR, we show that the C-terminus transitions from a rigid structure to a more dynamic structure as the solution is rendered acidic. We make previously unreported assignments of residues in the C-terminus of lipid embedded channels. Further, we also show evidence for hydrolysis of lipid head groups in proteoliposome samples during typical experimental timeframes.

## 1. Introduction

KcsA, the inward-rectifying potassium channel from the gram-positive soil bacterium *Streptomyces lividans*, has played a unique role in the structural biology of ion channels, being the first to be characterized in detail. The continuing progress on structural, biochemical, eletrophysiological, and biophysical characterization of KcsA makes it a uniquely rich model system for transmembrane transmembrane allosteric coupling [2,3] ion channel activation and inactivation mechanism [4,5], lipid-protein interactions [6,7] and virtually any other question concerning ion channels and membrane proteins. Yet, we do not yet have a full-length, atomic-resolution structure of KcsA in a lipid membrane. KcsA is composed of four identical subunits of 160 amino acids, each with two transmembrane spanning domains. While the transmembrane segments are relatively well characterized, the most mobile portions of KcsA, its extracellular termini in particular, present a unique set of challenges to resolve while in the membrane, despite their important roles.

### 1.2. Significance of the C-Terminus

The functionally crucial C-terminus is a case in point. The C-terminus of KcsA has been proposed to be to α-helix projecting perpendicular to the membrane[1,8–10] surface. Functionally, the C-terminus is important to the channel’s activation gating and pH dependence[9], as well as to its stability as a tetramer [11,12].

### 1.3. Lipid Interactions

Because activation of KcsA is lipid-dependent, details of the lipid environment are likely to be important for the dynamics of the loops and termini and for stabilizing conformations that are relevant for understanding function. KcsA has been shown to have an open-probability dependence on the lipid head group with the inner leaflet composition playing the key role[6]. The stability of the KcsA tetramer is also lipid dependent with anionic head groups (e.g. phosphatidic acid (PA), phosphoglycerol (PG), and phosphoserine (PS)) providing greater stability especially at low pH compared to lipids with net-neutral or net-positive headgroups [13]. Crystal structures [14–17], biochemical experiments[18], and NMR[19] have shown that KcsA routinely co-purifies with a diacyl lipid with a PG headgroup. Protein-lipid affinity experiments have shown that KcsA has a high affinity for the net-negatively charged lipids (PA, PE, and PG), with PG affinity being highest of all [20]. It is therefore likely that developing appropriate conditions for observation and characterization of the mobile termini by NMR or other methods will also depend on the lipids used. We therefore studied KcsA’s C-terminus in a biologically relevant, native-like lipid bilayer.

### 1.4. Prior NMR Studies

Many studies have applied high resolution cross polarization magic angle spinning solid state NMR (CPMAS) to make residue assignments to resonances of KcsA in proteoliposomes [21–25]. The protein residues that are detected and assigned are notably incomplete, emphasizing primarily the transmembrane helices (Figure 1).

**Figure 1.**
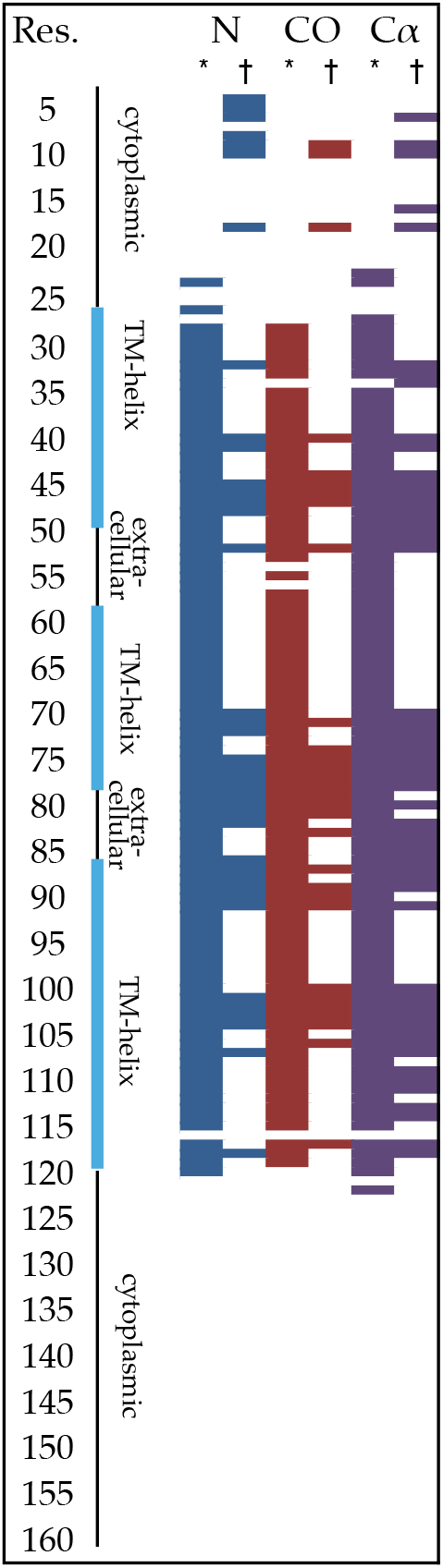
Solid-state NMR backbone assignments of KcsA in liposomes as described by the McDermott Group (*)[3, 23, 26, 27] or Baldus Group (†) [25, 28, 29]. Structural regions are indicated to the right of residue numbers.

Meanwhile, several efforts [5,30–33], most notably by Chill et al. [8], have provided nearly complete assignments for KcsA in detergent micelles. Solution studies have provided remarkable information about the dynamics [34] and secondary structure [8], and they have identified the major [35] and minor pH sensors [12]. Solution studies of KcsA often involve truncation mutants and high temperatures (e.g. > 45 °C). Specific studies of the excised C-terminus have also been conducted by solution NMR. The excised KcsA C-terminus (KcsA_112 – 160_) oligomerizes in solution, forming a tetramer as the pH decreases [12,31]. Tetramerization and pH dependence has been demonstrated for the C-terminus in full-length KcsA in MSP1D1 lipid nanodisks [36] with short-chained, unsaturated lipid 1,2-dimyristoyl-sn-glycero-3-phosphocholine (DMPC), with one tetrameric channels per disk. Using ^1^H-^15^N and ^1^H-^13^C TROSY data of ^15^N and methyl-labeled KcsA with an otherwise deuterated background, the authors identified two major states: a low pH state with 35 sharp amide resonances; and a second state at neutral pH that revolves only six broad resonances. These former studies in aggregate provide motivation to study the C-terminus of KcsA by NMR, potentially including hybrid solid state solution state NMR methods. Arguably, proteoliposomes provide a gold standard for authentic membrane environments. With that perspective in mind, we show that HR-MAS can be utilized to probe the C-terminus of KcsA while embedded in a membrane.

## 2. Results

Since the C-terminus of KcsA is not resolved by cross polarization magic angle spinning NMR (CP-MAS) nor by crystallography, we reasoned that the order and rigidity required by these methods is likely lacking. Therefore, we pursued strategies to detect these signals that are akin to liquid state NMR, including *J*-based NMR measurements which work well on isotropic systems. The liposome environment increases the tumbling time of embedded molecules and thus increases the anisotropy, and so the transmembrane portions of the proteins are not expected to be resolved in *J*-based experiments. On the other hand, proton-detected, *J*-based, high-resolution magic angle spinning NMR (HR-MAS) has unique potential to be useful to selectively detect portions that are significantly mobile (e.g., the termini) despite being anchored to a relatively immobile transmembrane domain.

### 2.1. HR-MAS Selectively Detects Signals from the KcsA C-Terminus

The ^1^H–^13^C HSQC HR-MAS spectrum of full-length KcsA was compared to a C-terminal truncation construct (KcsA-Δ125) both in 9:1 DOPE-DOPS liposomes (SI Figure 1). Protein-free liposomes were also examined to confirm the resonances arising from lipids. From this comparison, we concluded that much of the protein signal in spectra of the full-length construct arise from the C-terminus. By contrast, the relatively few signals observed in the KcsA-Δ125 sample arise mainly from the synthetic phospholipids into which the protein is reconstituted. Similarly, ^1^H-^15^N HSQC spectra (Figure 2) also show that most (though not all) resonances are eliminated when the C-terminus is truncated, indicating that HR-MAS signals of KcsA principally arise from the C-terminus.

**Figure 2.**
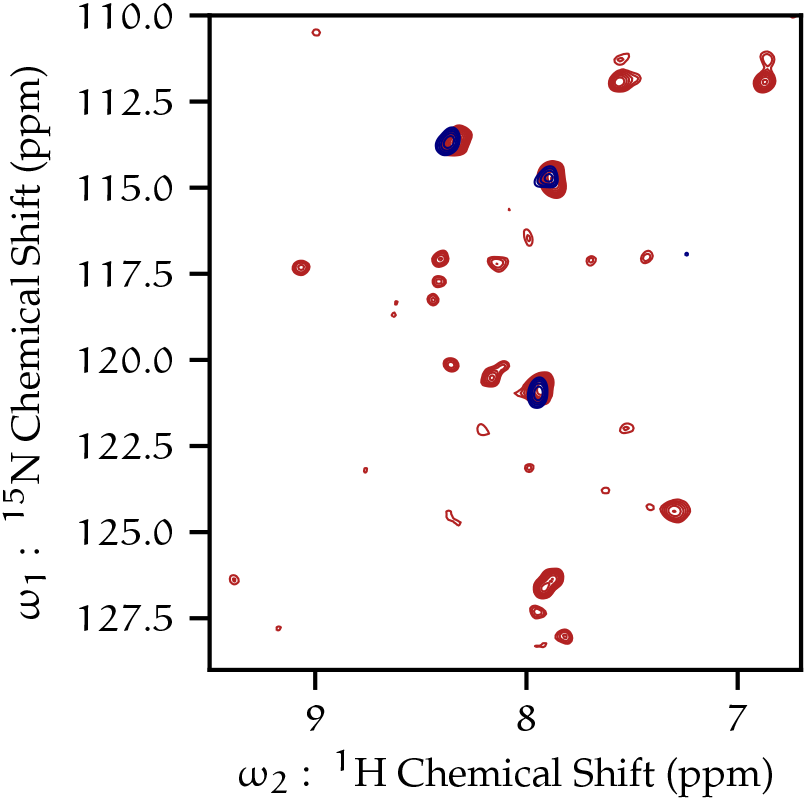
^1^H–^15^N HSQC by HR-MAS of full-length wild-type KcsA (red), and KcsA-Δ125 (navy); both samples pH 7.25, 50 mM K^+^ 308 K, 5 kHz MAS.

### 2.2. Cleavage of the KcsA C-terminus causes conformational heterogenity near selectivity filter

To inspect for changes to the KcsA transmembrane domain caused by cleavage of its C-terminus CP-MAS ^13^C–^13^C proton driven spin diffusion (DARR) spectrum of KcsA was collected of KcsA-Δ125 in liposomes at neutral pH and compared to the full-length construct of KcsA (SI Figure 3). Previous work shows the ^13^C–^13^C spectrum of full-length KcsA (FL-KcsA) in liposomes records resonances arising from KcsA’s transmembrane domain [23], therefore SI Figure 3 shows that the transmembrane domain of KcsA-Δ125 is well-folded and ^13^C enriched. However, the KcsA-FL sample has several additional resonances that KcsA-Δ125 sample is missing. Using chemical shift assignments from previous studies in our group [23] were used to assign the resonances that are present in KcsA-FL but absent in KcsA-Δ125. Signal from residues I38, L59, T74, D80, Y82, T85 is absent in the spectrum of the truncated construct and all of these residues are located in either the extracellular loop domains or within the selectivity filter (Figure 3), with the exception of I38.

**Figure 3.**
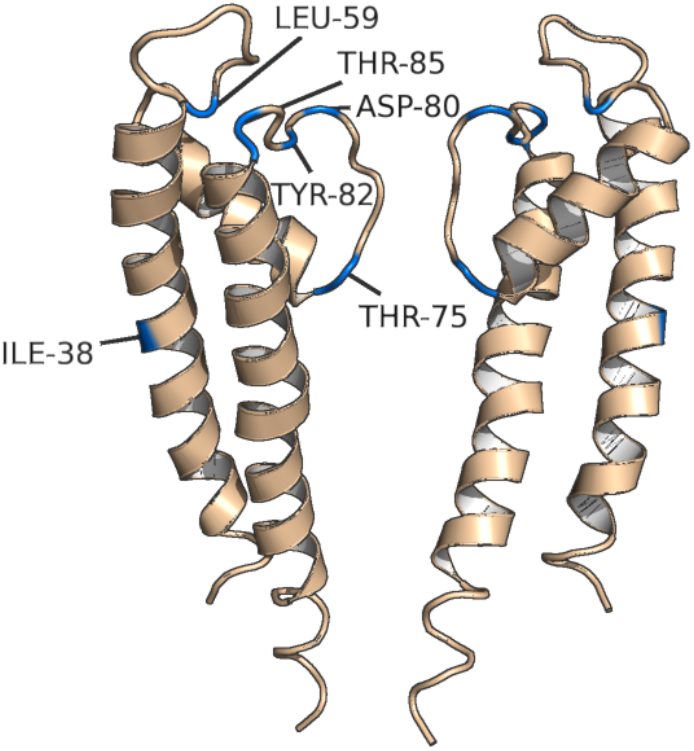
Resonances missing from CP-MAS ^13^C–^13^C spectrum of KcsA–Δ125 (blue) as compared to fulllength KcsA mapped onto PDB: 1BL8 [37] visualized with PyMol, two of four identical subunits are hidden for clarity.

Previous work in our group has proven T74CA-CB chemical shift to be an indicator of the K^+^ apo and bound states of the channel [3,26]. These previous studies show perturbations in the chemical shifts, not absence of resonances. The first crystal structure of KcsA was of a KcsA-Δ125 construct. The resonances that are present in KcsA-FL and absent in KcsA-Δ125 are mapped onto that crystal structure of a KcsA-Δ125 construct from Doyle et al. [37] in Figure 3. The most likely explanation of these NMR data, however, is that the C-terminal truncation leads to much greater dynamics, leading to conformational heterogeneity of the regions at the interface between KcsA’s loop regions and selectivity filter. Previous work has shown that deletion of the C-terminus (KcsA-Δ125), impairs the ability for the protein to assemble into a tetramer [11,38] and slightly decreased open-probability [6].

### 2.3. Low pH leads to more dynamics of the C-terminus of KcsA embedded into liposomes

Although the ^1^H-^15^N HSQC HR-MAS spectrum of KcsA in liposomes at neutral pH contains few resonances, most of which originate from the C-terminus, by contrast, when the pH of the sample is lowered to pH 4, both the number and the distribution of resonances in the spectrum increases (Figure 4). These spectra were collected on a fractionally deuterated protein sample. Analogous phenomena were observed in fully protonated samples as well (though those spectra are less resolved, data not shown).

**Figure 4.**
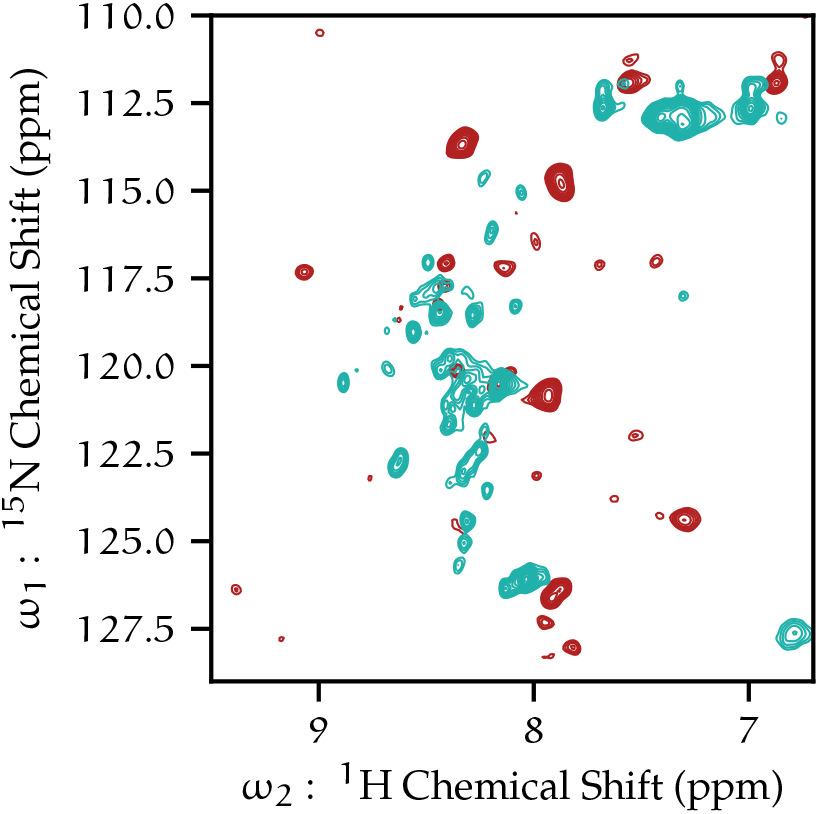
^1^H-^15^N HSQC of KcsA at pH 7.25 (red), and F-^2^H-KcsA at pH 4.0 (blue) by HR-MAS. 5 kHz MAS, 308 K, 50 mM K+.

The dramatic increase in the number of resolved resonances and an overall increase in signal strength suggests that KcsA is undergoing much greater conformational dynamics at low pH. This would lead to greater averaging of orientation-dependent dipolar couplings and chemical shift anisotropy, effectively increasing the transverse-relaxation time (*T*_2_), leading to increased transfer efficiency in the *J*-based HSQC experiment.

### 2.4. Leucine as a C-terminal Conformational Indicator

The conformational and dynamical changes the protein undergoes from neutral to low pH can also be observed in the ^1^H–^13^C HSQC. One reliable identifier of pH-induced state is the leucine ^1^H-^13^Cδ (methyl) correlations in the HSQC (as identified by chemical shift and TOCSY fingerprint). The peaks show reversible shifts when pH drops, reflecting systematic changes to the entire spectrum, and to the pair of leucine in particular (Figure 5). These leucine resonances are not present in the C-terminal truncation construct, confirming these resonances arise from the C-terminus, which has a total of three leucines (L144, L151, L155). The difference in the shift dispersion is a strong indication that at low pH the conformation of the C-terminus is more heterogeneous than the conformation at neutral pH where the channel is expected to be in the closed conformation.

**Figure 5.**
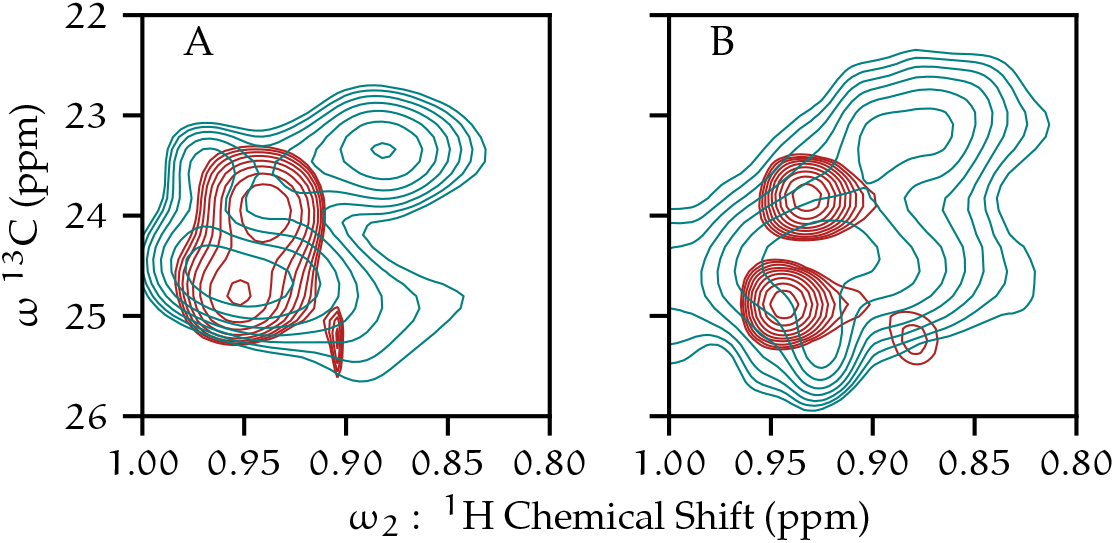
1H–13C HSQC spectra of leucine Hδ of KcsA in liposomes. pH 7.25 (red contours) and pH 4.0 (blue contours) are shown, with fully protonated KcsA samples (A) and fractionally deuterated KcsA samples (B).

**Figure 6.**
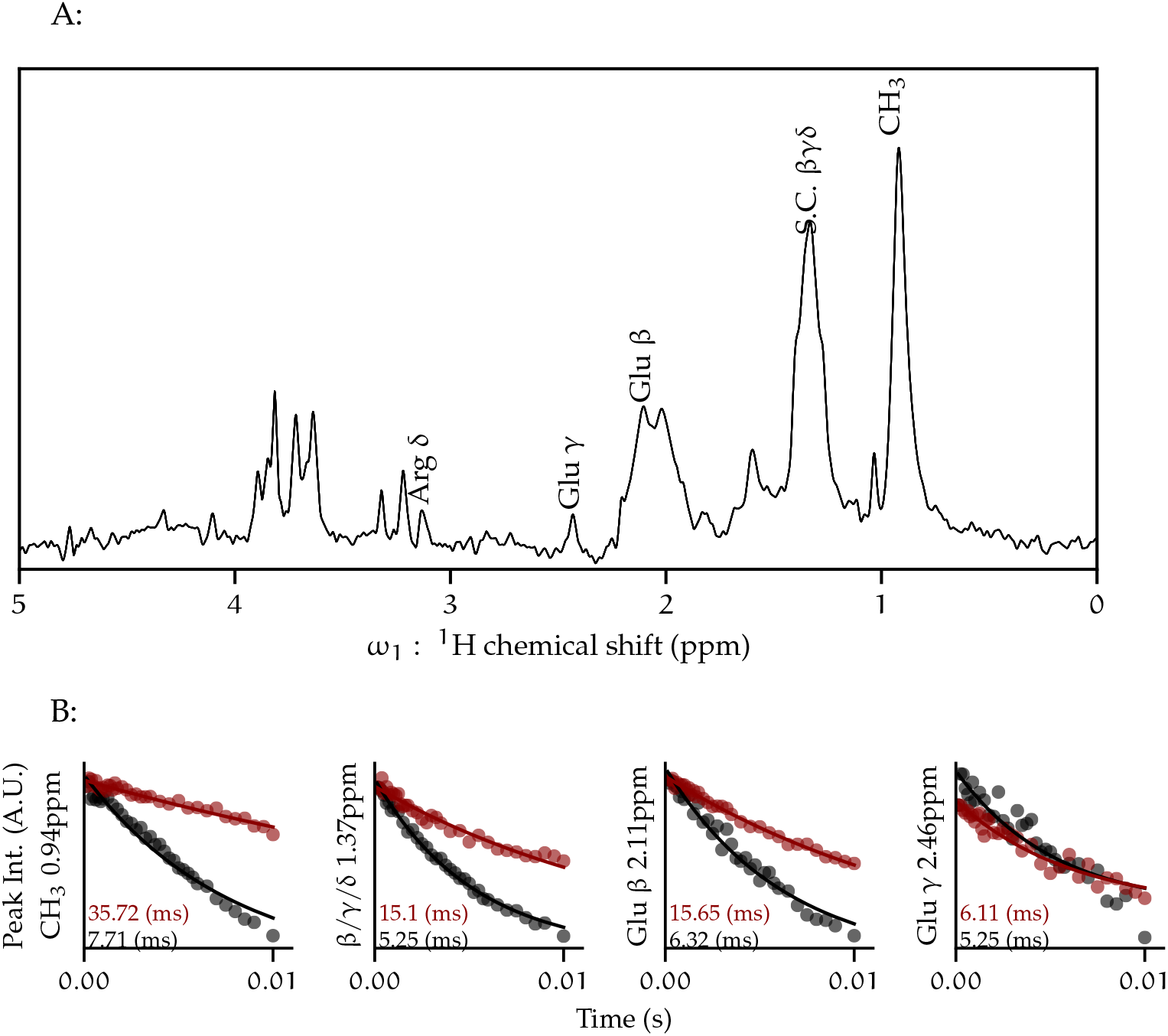
The *T*_2_ relaxation time is lengthened due to fractional deuteration of KcsA. (A) ^1^H-^13^C double-INEPT (1D refocused HSQC) spectrum of U-^1^H, ^13^C, ^15^N-KcsA in proteoliposomes (9:1 DOPE-DOPS) at pH 4.0, 50 mM K^+^, with peak annotations. Assignments are based on 2D and 3D data. 9 kHz MAS, 308 K. (B) ^13^C relaxation of selected peaks from 1D double-INEPT for fully protonated KcsA (black), and fractionally deuterated KcsA (red). Both samples U-^13^C,^15^N at pH 4, 50mM K^+^.

**Figure 7.**
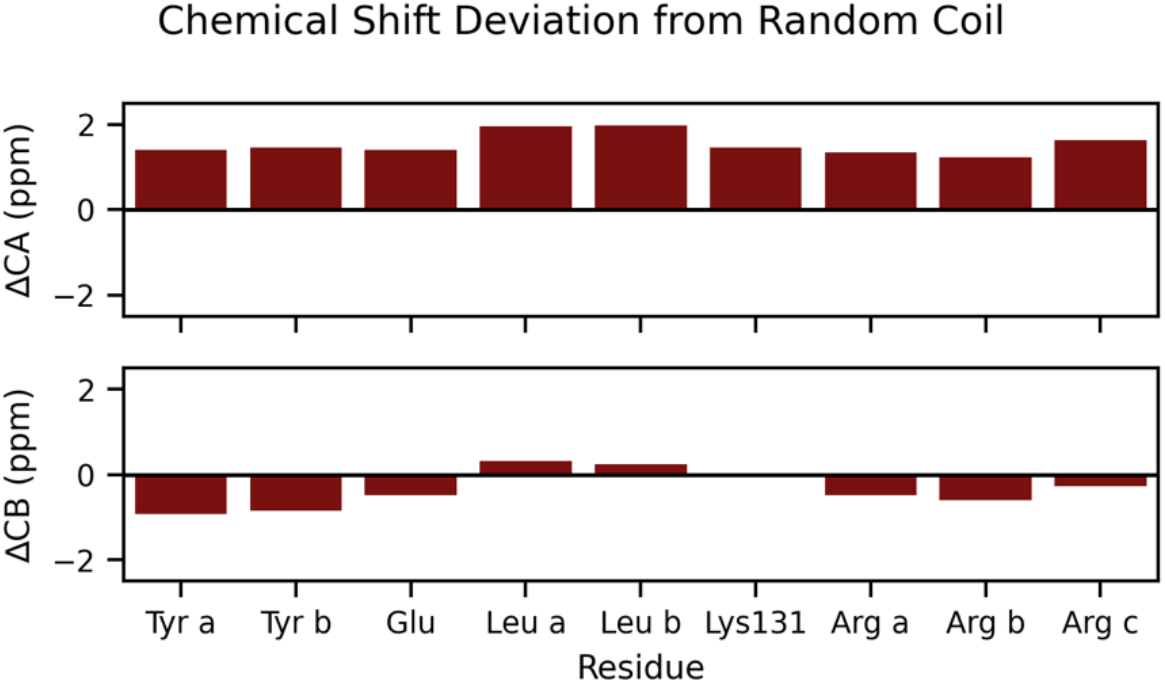
Chemical shift deviations from random coil of residues in the KcsA C-terminus at Low pH condition KcsA. From hCCH-TOCSY data, U-^1^H, ^13^C, ^15^N-KcsA in proteoliposomes (9:1 DOPE-DOPS) at pH 7.2, 50 mM K^+^.

### 2.5. ^13^C T_2_ Relaxation

Site-specific ^13^C transverse relaxation of protein signal were measured to optimize polarization transfer by collecting a gradient-selected pep-sensitized HSQC with a variable length dephasing period before *T*_1_ evolution (SI Table 1: ^13^C *T*_2_s). The average protein ^13^C *T*_2_ relaxation rate was 2.9 ms and the average for lipid headgroup ^13^C was 5.2 ms for the protonated sample at a sample temperature of 308K, 9 kHz MAS, and 10 kHz hetero-nuclear decoupling. The optimal transfer time for a single Cα-N polarization transfer for the refocused-INEPT (in the absence of relaxation) is 1/(2 *J*), and since *J*_N-Cα_ is typically 7-11 Hz [39], the optimal transfer time would be expected to be 45 to 70 ms. However accounting for relaxation with measured *T*_2_ values the optimal transfer is 2.9 ms, with a maximum theoretical signal yield of only 3.7 % when compared to yield possible in the absence of relaxation, as predicted from the density function of the refocused INEPT experiment, 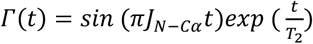 [40]. Under these conditions even the highest sensitivity ^1^H-detected backbone experiments, such as the HNCaH and HNCO (out-and-back), which require multiple INEPT transfers are not feasible with fully protonated samples.

### 2.6. Fractional Deuteration

Anisotropic interactions, particularly dipolar couplings, lead to shorter *T*_2_ relaxation values for proteins. Residual dipolar couplings with ^1^H nuclei vastly attenuate *J*-based coherence transfer in backbone experiments. Replacing ^1^H with ^2^H in proteins is a well-established method of reducing these dipolar couplings, thus increasing *T*_2_ times, improving resolution and coherence transfer efficiency [41,42]. Although reports of highly- [5,33,36] and perdeuterated [8] KcsA exist in the literature, expression of perdeuterated KcsA led to poor yields in our hands. Instead, two different fractional deuteration schemes were successful. Previous work has found that expressing proteins in D2O minimal media supplemented with U-^1^H, ^13^C-glucose and ^15^N-ammonium chloride leads to proteins that have very high levels of deuteration at the H-Cα position and fractional deuteration in the sidechains that vary by amino acid [43]. Here, we expressed KcsA in this manner, and incorporated it into 9:1 DOPE-DOPS liposomes. ^13^C *T*_2_ relaxation profiles were determined using a 1D back-INEPT experiment with a dephasing period. In addition to profiles for the methyl region and the aliphatic region, Glu-^13^Cβ and Glu-^13^C γ profiles are displayed because they are easily identified on the 1D spectrum of both samples. The results show that the relaxation profile is vastly improved of the F-^2^H sample over the uniformly ^1^H labeled sample. The Glu-Cγ profiles are similar for both samples, while agrees well with previous observations that both glutamate γ protons tend to remain protonated in this labeling scheme and a portion of the Glu-Hβ remain protonated in this labeling scheme [28]. The real testament to the effectiveness of fractional deuteration is that 3D correlations were possible only with deuteration. With the fully protonated sample, signal was insufficient to collect the 2D HNCA or HNCO correlations experiments, while with F-^2^H-KcsA, full 3D datasets were obtained showing 13 well resolved resonances (SI Figures 8 and 9).

### 2.7. Chemical Shift Data

### 2.8. Lipid Stability in NMR Samples

Solid-state NMR samples routinely spend a week or more in the probe. Near complete assignments (except highly degenerate resonances in the aliphatic chain) of KcsA proteoliposome lipids could be made in ^1^H–^13^C HSQC HR-MAS data (Table 2). At least six resonances were identified as appeared frequently in KcsA proteoliposome data that do not correspond to known lipid species or are consistent with protein chemical shifts. hCCH-TOCSY experiments (SI Figure 7), reveal spin systems that correspond to the literature values of free glycerol [44,45] and free ethanolamine [45]. Since neither glycerol nor ethanolamine is used in the preparation in these samples, and the signal strength from these compounds is too high to suggest trace contaminant the signal and chemical logic, both suggest that these components result from degradation of the lipids. The chemical shifts of the products and their presence in various samples is further documented in Table 2 with additional detail in SI Table 2. We and others have previously reported that exogenous lipids co-purify with KcsA expressed in *E. coli* containing a phosphoglycerol head group [19]. The copurified lipids can be identified because they are expected (like the purified protein) to have uniform enrichment of ^13^C and ^15^N, in contrast to exogenously lipids added during reconstitution which would have natural abundant ^13^C and ^15^N content. The most plausible source of ethanolamine is from hydrolyzed exogenous PE lipid headgroups. We have not identified the catalyst for this hydrolysis. The HSQC signal for free glycerol is invariably much greater than that of free ethanolamine (when ethanolamine is detectable). This suggests to us that at least some of the glycerol arises from hydrolysis of co-purifying PG with the ^13^C-enrichment explaining higher signal despite the much lower concentration of PG lipids as compared with PE. Phosphoglycerol or free glycerol is present in nearly all KcsA samples, with most containing signal from both. The strong ^13^C-^13^C correlations in hCCH-TOCSY data suggests that at least a portion of this glycerol signal is isotopically enriched (SI Figure 7) suggesting the co-purifying PG lipid is responsible in part for the glycerol. However, the intense signal from glycerol in ^1^H direct excitation spectra (Figure 8) demonstrates the glycerol is at a concentration much greater than is possible from hydrolyzed PG alone.

**Table 1.** Amino acid types identified in hCCH-TOCSY experiment of KcsA in 9 : 1 DOPE-DOPS at pH 7.2

| Residue | Cα | Cβ | Cγ | Cδ1 | Cδ2 | Cζ | Hα | Hβ1 | Hβ2 | Hγ | Hδ |
| --- | --- | --- | --- | --- | --- | --- | --- | --- | --- | --- | --- |
| Lys 131 | 57.5 | 32.6 |  |  |  | 42.1 | 3.83 |  |  |  |  |
| Arg a | 57.1 | 30.4 | 25.6 | 41.8 |  |  | 3.86 | 2.09 |  | 1.63 | 3.15 |
| Arg b | 57.0 | 30.2 | 25.6 | 41.8 |  |  | 3.83 | 2.06 |  | 1.63 | 3.14 |
| Arg c | 57.4 | 30.6 | 27.8 | 42.1 |  |  | 3.77 | 1.96 |  | 1.62 | 3.14 |
| Leu a | 56.3 | 42.7 |  | 23.7 | 24.9 |  | 3.81 | 1.82 | 1.76 |  |  |
| Leu b | 56.3 | 42.6 |  | 23.8 | 24.8 |  | 3.81 | 1.82 | 1.76 | 1.82 |  |
| Phe/Tyr a | 58.9 | 38.3 |  |  |  |  | 4.01 | 3.28 |  |  |  |
| Phe/Tyr b | 58.9 | 38.4 |  |  |  |  | 4.02 | 3.13 |  |  |  |
| Glu a | 57.5 | 29.7 | 36.4 |  |  |  | 3.83 | 2.12 |  | 2.44 |  |
| Glu b | 57.1 | 29.6 | 36.4 |  |  |  | 4.08 | 2.20 |  | 2.44 |  |

**Table 2.** Chemical shift assignments for liposome lipids and lipid hydrolysis products measured in this paper at pH 4-7, 50 mM KCl, 308K, 5 kHz MAS. See Figure 8 for nomenclature.

| Site | <sup>1</sup>H (ppm) | STD | <sup>13</sup>C (ppm) | STD |
| --- | --- | --- | --- | --- |
| Fatty acid 12 ($\omega$) | 0.88 | 0.018 | 16.6 | 0.12 |
| Fatty acid 11 ($\omega - 1$) | 1.3 | 0.017 | 25.3 | 0.06 |
| Fatty acid 8 ($\text{CH}_2\text{-HC=C}$) | 2.02 | 0.038 | 29.9 | 0.19 |
| Fatty acid 6 ($\text{CH}_2$) | 1.29 | 0.022 | 32.1 | 0.1 |
| Fatty acid 10 ($\omega - 2$) | 1.27 | 0.02 | 34.6 | 0.08 |
| Fatty acid 4 ($\alpha\text{-CH}_2$) | 2.33 | 0.033 | 36.6 | 0.24 |
| PE $\beta$ | 3.24 | 0.037 | 43.3 | 0.37 |
| Ethanolamine $\beta$ | 3.19 | 0.088 | 44.2 | 0.1 |
| Ethanolamine $\alpha$ | 3.84 | 0.095 | 60.3 | 0.2 |
| PE $\alpha$ | 4.11 | 0.027 | 64.6 | 0.48 |
| PG $\gamma$ + glycerol a | 3.67 | 0.025 | 65.4 | 0.1 |
| PG $\gamma$ + glycerol b | 3.61 | 0.044 | 65.4 | 0.09 |
| G3 a | 4.23 | 0.028 | 67 | 2.2 |
| G3 b | 4.46 | 0.027 | 67 | 2.3 |
| G1 | 4.3 | 0.23 | 67.5 | 3.4 |
| PG $\alpha$ a | 3.94 | 0.022 | 70 | 1.8 |
| PG $\alpha$ b | 3.88 | 0.025 | 70 | 1.8 |
| PG $\beta$ | 3.93 | 0.028 | 72 | 2.2 |
| G2 | 5.26 | 0.012 | 73.3 | 0.15 |
| glycerol ($\text{CH}_1$) | 3.83 | 0.035 | 74 | 2.4 |
| Fatty Acid 9 ($\text{HC=C}$) | 5.31 | 0.027 | 132.2 | 0.12 |

**Figure 8.**
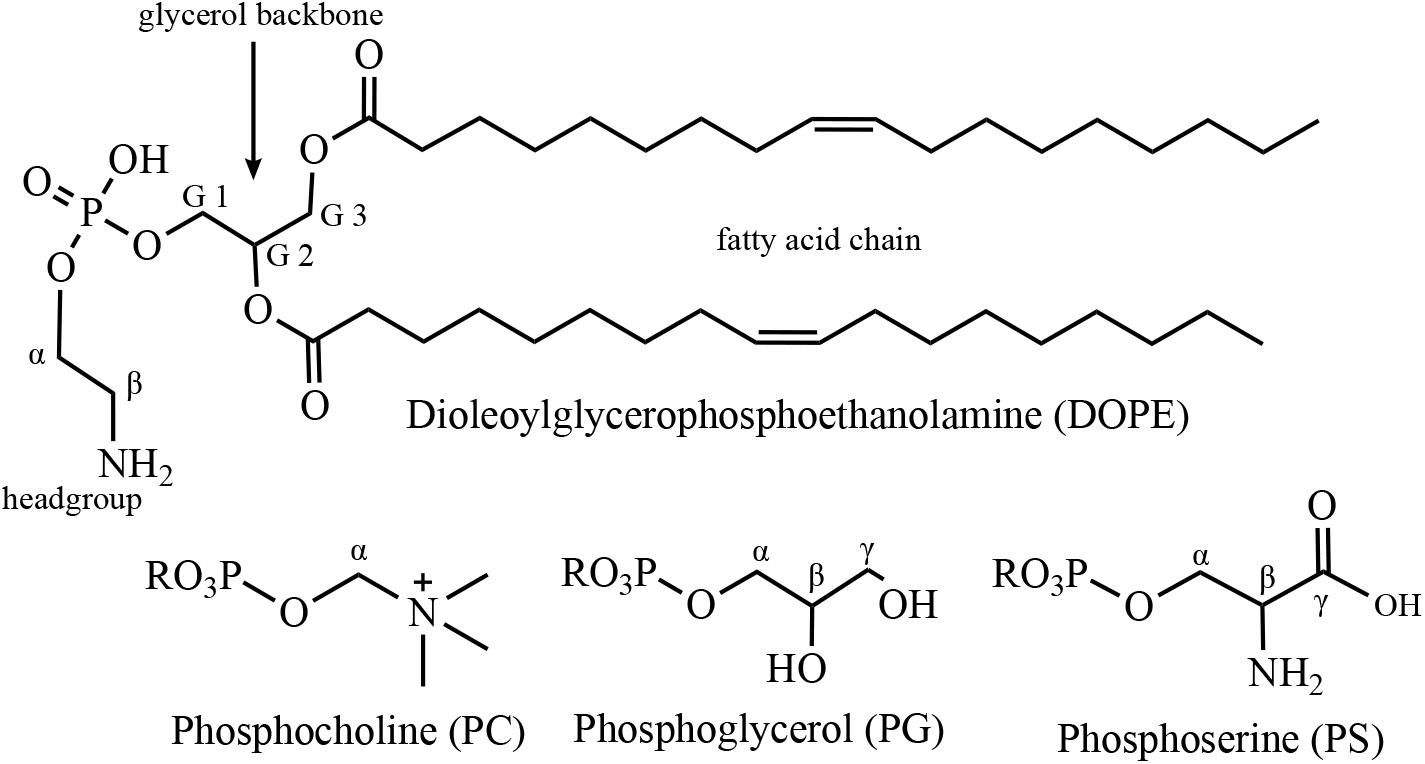
Structure of lipids of interest to this study and their nomenclature

In Figure 8 we demonstrate that a direct polarization ^1^H experiment with pre-saturation conducted at 308K in an MAS probe is sufficient to diagnose the presence of lipid hydrolysis products. The experiments shown are for an HR-MAS probe, however we find this experiment can be accomplished effectively in an ^1^H/^13^C/^15^N e-free CP-MAS probe configured for ^1^H-dectection. The sharp peaks arising from mobile solutes of lipid hydrolysis were identified by chemical shift and confirmed through ^1^H-^13^C HSQC and hCCH-TOCSY data (see SI Figure 4 for example assignment data). The principal small molecule (sharp peaked) signal are from buffer components and water soluble hydrolysis products of lipids such as glycerol and ethanol amine.

We took a survey of preserved samples from previous studies of KcsA liposomes from our group and for each sample investigated, a gradient-selected, phase sensitive ^1^H-^13^C HSQC by HR-MAS with the sample at 308 K and 5 kHz MAS was collected. The spectra were examined for the presence of PG, ethanolamine, and glycerol, and the chemical shifts of these compounds were then used to assign peaks in quantitative ^1^H spectra of the same samples. The results are summarized in SI Table 2. The shifts from HSQC data were used to assign the peaks in the ^1^H data, and thereby the presence or absence of the glycerol and ethanol amine was verified.

### 3.8. Effect of MAS Centrifugation on Samples

To homogenize the sample and create ULVs, KcsA proteoliposomes were pelleted following dialysis and then subjected to 25 rounds of freeze-thaw using liquid nitrogen and a 30° C water bath, leading to the formation of proteoliposomes of unilamellar as well as multilamellar or oligomellar structures (SI Figure 6), similar to previous reports [46,47].

To characterize the proteoliposomes resulting from this freeze thaw procedure, we loaded the samples on to a uniform sucrose and buffer gradient (5-60 %) and performed isopycnic ultracentrifugation. In order to visualize the proteloposomes on the column and quantify the lipid concentration, the we included a rhoadamine-congated lipid (1,2-dioleoyl-sn-glycero-3-phosphoethanolamine-N-(lissamine rhodamine B sulfonyl) (Rhod PE) in the lipid mixture. The lipid mixture had a mass ratio of 900:10:1 PE-PS-Rhod PE and total lipid to protein mass ratio was 1:1.

Following centriguation, aliquots were taken, and portions of each aliquot were used to determine protein concentration by solubilizing proteliposomes with a solution of Triton-X100 and bromophenyl blue and comparing UV-VIS absorbance at 610 nm with a standard curve [48] and total lipids were extracted from aliquots using chloroform and methanol [49] and quantified using a standard curve based on UV-VIS absorbance from Rhod-PE at 560 nm. Density of aliquots was determined by analytical balance and calibrated micropipette using low retention tips.

The isopycnic gradient experiment reveals to distinct sets of populations distinguished by their densities (Table 3). All of the data described in this paper (and many others) would presumably contain both of these populations as we were unable to efficiently separate these populations.

**Table 3.** Fractions of KcsA and DOPE–DOPS–Rhod-PE proteoliposomes from isopycnic sucrose gradient ultracentrifugation.

| Aliquot Density (g / cm<sup>3</sup>) | Lipid : Protein | Proportion of loaded protein (%) |
| --- | --- | --- |
| 1.02 ± 0.01 | 7 ± 1 | 20 ± 10 |
| 1.07 ± 0.01 | 0.8 ± 0.1 | 70 ± 10 |

We hypothesize that we could mitigate the effect of centrifugal force caused by magic angle spinning by density matching our samples to the buffer matrix in which they were suspended. We hypothesized that the placing the liposomes in a density matched matrix will diminish the deleterious mechanical effects of magic angle spinning without affecting the spectroscopic benefits. However, despite many repeated attempts at isopycnic preperations, we were unable to achieve total protein concentrations of greater than 5% in aliquots. As NMR is a relatively insensitive technique, and therefore sample concentration a critical feature of obtaining strong signal in a reasonable amount of time, we sought alternative methods to density match the sample.

MAS can be expected to exert pressure on the proteoliposomes, potentially changing their morphology or hydration, and therefore affect both function and NMR detection. To reduce net forces on the proteoliposomes, we sought to prepare an isopycnic solution of the proteoliposomes. To roughly match the density of the KcsA pellet to the buffer, we ‘floated’ the unilamellar proteoliposomes pellet by adding aliquots of 60 % w/w sucrose-augmented buffer to a tube containing the pellet, then vortexed and centrifuged at 21,130*g* in a bench top centrifuge and observed whether the pellet sank or floated. It was found that this process can be reversed (i.e. cause a floating pellet to sink) by adding an aliquot of buffer without sucrose to the tube. The process can be repeated in either direction indefinitely. Thus, this experiment was dubbed a reversible ‘elevator’ experiment and provided a means to closely titrating the density to match KcsA proteoliposomes. Because we used transparent rotor inserts, we were able to verify the floating pellet does indeed migrate to the center of the rotor during MAS (Figure 9).

**Figure 9.**
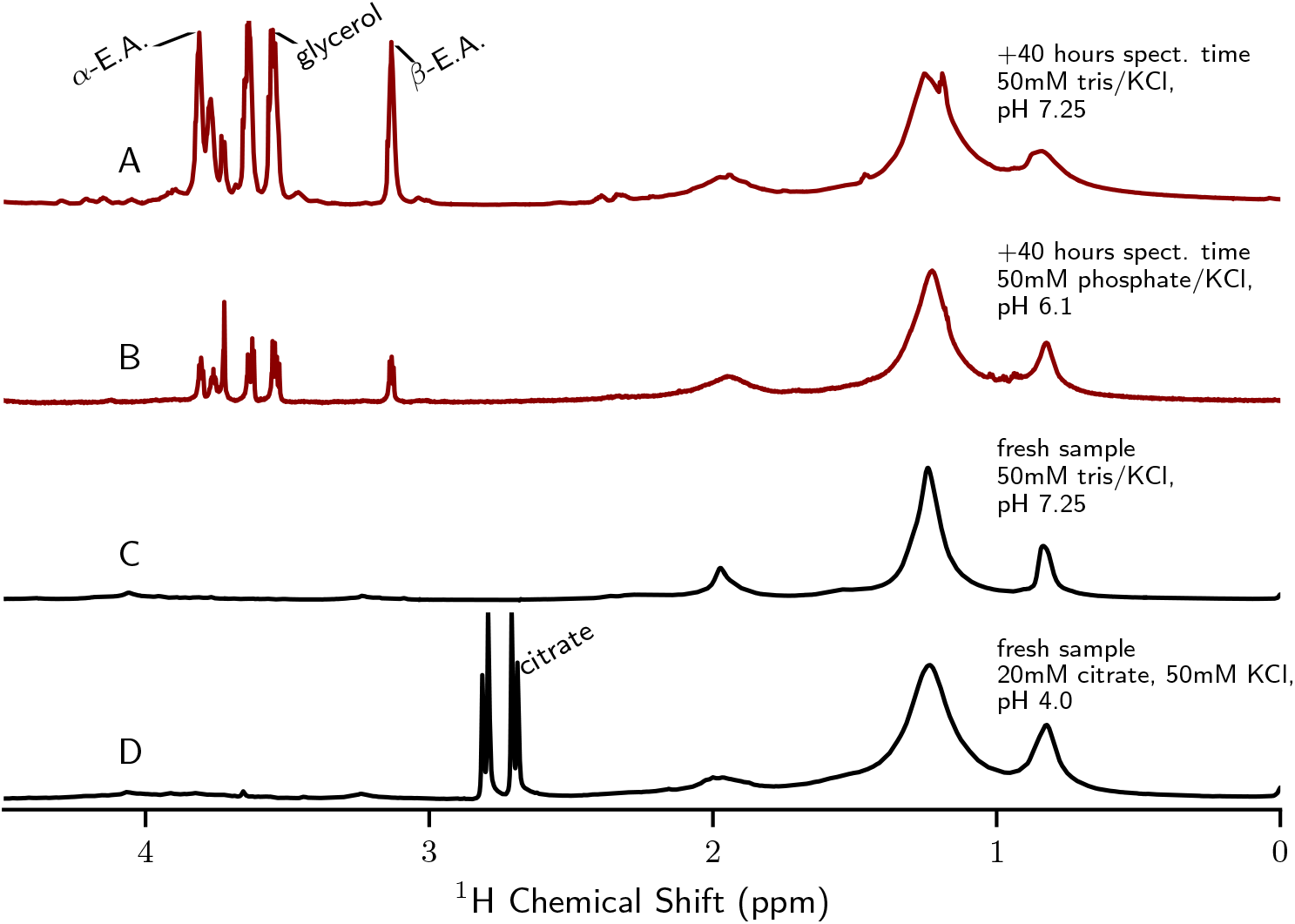
Quantitative ^1^H direct excitation MAS spectra (with water presaturation) of KcsA proteoliposomes samples. Samples that contained free ethanolamine signal are displayed in red. The samples that contain only PE (intact lipid) and no ethanolamine are displayed in black. All spectra were collected at 308 K, 5 kHz MAS. Spectra are normalized to the bulk CH2 signal (~1.2 ppm). All samples are U-^1^H-^13^C-^15^N-KcsA in 9:1 DOPE liposomes, LPR = 1. Free glycerol, ethanolamine (E.A.), and citrate peaks are labeled.

**Figure 10.**
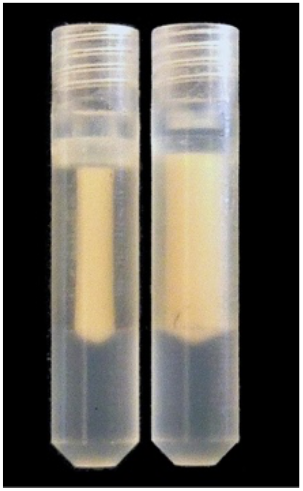
KcsA proteoliposomes in MAS rotor inserts post-experiment. Left: ‘Floating’ sample in sucrose-augmented buffer, and Right: pelleted sample with no sucrose in buffer.

We did not identify dramatic differences in ^1^H–^13^C or ^1^H–^15^N HSQC by HR-MAS spectra of KcsA unilamellar proteoliposomes that were ‘floating’ versus pelleted (data not shown). Cryo-electron microscopy (SI Figure 6) shows that before MAS, many small unilamellar vesicles are present and following MAS, there are no unilamellar vesicles that could be found. Instead, post-MAS of pelleted and floating samples are less well ordered and show multilamellar, oligolamellar, and aggregated morphology. This suggests that the floating samples are not protected from changes to lipid morphology under MAS conditions.

## 3. Discussion

Here we present evidence that KcsA in model liposomes assumes distinct conformations at pH 7.25, where the channel is inactive, as compared with at pH 4.0, under which conditions the channels is able to activate. The specific lipid model we have selected here has been shown to be compatible with activity including activation, inactivation [22] and allosteric coupling between sites within KcsA [3,23], and thus these sample conditions have been appropriate for studies that connect structure to channel function.

The conformational switch presented here is consistent with previously proposed structural models. A sequential spin-labeled, ESR structure of KcsA in bilayers indicated significant mobility of the C-terminus at pH 4 [9]. On the other hand, full-length KcsA with its C-terminus stabilized by Fab antibodies at neutral pH was rigid enough to capture by X-ray crystallography [1], suggesting that without the antibodies the C-terminus was highly mobile.

This study and others [36,50] reports two distinct dynamical regimes of the KcsA C-terminus as a function of pH. Dramatic changes can be observed in the ^1^H–^15^N HSQC data (SI Figure 2), with many more peaks with much narrower line widths at low pH than compared to neutral pH. This is particularly clear when comparing fractionally deuterated samples but is also observed with fully protonated samples, showing reproducibility and independence of the labeling scheme. These data indicate that as the pH is lowered the KcsA the C-terminus moves more freely.

Our analysis of hCCH-TOCSY spectra showed ten resolved groups of resonances that can be assigned by amino acid type (summarized in Table 1). These data alone established an additional new site-specific spectral assignment (Lys-131). We also identify three arginine, two leucine, two aromatic (Phe or Tyr), and glutamic acid spin systems. We are unable to definitively distinguish between Phe/Tyr though Tyr is more likely than the respective alternative based on canonical chemical shift data. All of these spin systems absent in the spectrum of KcsA-Δ125. So, the resonances are in almost certainly from the C-terminus. Previous x-ray structure of full-length KcsA in the closed confirmation [1] and whose data agrees with EPR structural data [9] suggest that, in the C-terminus, Phe-125, Arg-127, Lys-131, Glu-135, Arg-139, Arg-142, Glu-146, Arg-159, Arg-160 are solvent exposed and are therefore most likely to have resolved side-chain resonances. The most compact sequence of amino acid that conforms to the evidence would be from Phe-125 to Leu-151 (<u>F</u>V<u>R</u>HS<u>EK</u>AA<u>EE</u>A<u>Y</u>T<u>R</u>TT<u>R</u>A<u>L</u>H<u>ER</u>FD<u>RL</u>), and even within this short sequence several arginine and glutamic acid residues would be unresolved amongst amino acids type-resolved. According to the previous structural models, all of the candidate sites for hydrophilic residues (Lys, Arg X3, Glu X2) are on the solvent-exposed surface, whereas the leucine side chains are likely to be facing other protein subunits, and the phenylalanine and tyrosine

Through hCCH-TOCSY data and selective cleavage of KcsA, we have identified several residues by type that reside in the C-terminus at neutral pH. The ^13^Cα resonances of these residues have systematically higher shift values than mean values for random coil (Figure 7), suggesting that the residues we detect are likely to be in helical arrangement[51]. Several other studies have examined chemical shift deviations, finding KcsA typically has particularly elevated shift values (often > 6ppm) for KcsA tetramers in detergent micelles[8], and similar behavior in the C-terminal domain alone solubilized in water when it spontaneously tetramerizes at high concentration [31]. Both those studies, however also find small regions within the C-terminus at neutral pH that have less dramatic shift elevations (+ 1-2 ppm), which are putatively characterized as more dynamic, less rigid helices. Crystal structures [1,9] and EPR [52] measurements of KcsA in lipid mimics at neutral pH support further support the hypothesis of the C-terminus existing as series of linked helice. Our study, conducted in the lipid bilayer, comports with this structural model. Specifically, it is likely given the modest changes in shift, that the resonances we detect are from the more mobile portions and that the most rigid portions of the C-terminus in the tightest helices remain unresolved at our conditions.

This work also identified leucine ^1^H-^13^Cδ HSQC peaks that provide markers for the low versus high pH states (Figure 5). Specifically, we reproducibly observe a set of sharp leucine peaks at neutral pH and at low pH more peaks that are generally broader. This same behavior has been previously observed of the C-terminal domain alone solubilized in water [31]. EPR measurements in particular [52] suggesting L151 undergoes significant conformational rearrangement when the channel goes from a neutral pH to a low pH environment. That we resolve additional leucine resonances at low pH underscores our conclusions that the KcsA C-terminus becomes more mobile at low pH. Yet it is not clear why in our data, supported by soluble C-termini studies, that leucines in the C-terminus show the opposite trend of other resonances in becoming broader. These data cannot distinguish if the apparent line broadening is due to relaxation effects or due to conformational heterogeneity. Regardless, these findings support the use of leucine to distinguish the high pH and low pH conditions.

The data presented here show that HR-MAS is a viable method to investigate the structure of KcsA’s C-terminal domain. There are several interesting functional questions that could be addressed with this system. For example, depleting the system of K^+^ in the low pH state leads to structural changes and subsequently to the inactivation of KcsA, with a strong allosteric connection between binding of potassium at the selectivity filter and protonation of pH sensors at E118 and E120 [3]. HR-MAS could detect whether the C-terminus also has an allosterically induced conformational change, for example reverting to the more rigid structure upon depletion of K^+^ from the selectivity filter.

### 4.1. Lipid Models

In these pages we demonstrate that ^1^H-dected NMR can be conducted on KcsA embedded in proteoliposomes, a highly cell-like model system. Whereas previous ^1^H-detected studies of KcsA have been performed in model systems somewhat more removed from a native-like environment. Lipid-protein nanodisks have shown promise for the study of KcsA. Work on KcsA in PC nanodisks showed that the C-terminus has a pH dependent conformation, which this work supports in 9 : 1 DOPE-DOPS liposomes [36]. This suggests that the lipid milieu does not determine the C-terminal helix bundle dissociation at low pH. One suggested function of the C-terminus helix bundling is that the contacts made across monomers may aid the stabilization of the KcsA tetramer at neutral pH [50]. Those data were collected on a solution-NMR instrument and found that resonances were too broad and the *T*_2_ values too short to conduct 3D backbone experiments. Here, we have demonstrated the ability to conduct 3D experiments in liposomes, and we might see significant increases in resolution and improvements in *T*_2_ relaxation by combining both nanodisks and HR-MAS. Here, proteoliposomes are not expected to be rapidly tumbling. Instead, the protein diffusion through the lipid bilayer is the largest source of translational motion for the entire protein, and then individual domains such as the C-terminus, are likely to undergo further localized movement that can decrease anisotropic interactions. Lipid composition varies the rate at which lipids and proteins diffuse through the membrane surface. The lipid composition and the protein-to-lipid ratio could be tuned to produce more rapid diffusion and which might lead to improved relaxation and resolution characteristics. Lateral diffusion of lipid probes is inversely proportional to bilayer thickness [53]. Lipid hydration is very important in determining diffusion, with maximum diffusion rates occurring above 40 % water by mass[54]. Lipid acyl chain composition changes the diffusion rates, with DPPC (saturated lipid with 16 carbon chain), diffusing at more than two times the rate of DOPC (single cis-double bond per chain and 18 carbon chains) [55]. Headgroups play a dramatic role as well, with DOPG diffusing at twice the rate of DOPC [53]. Most of these studies named here are examining a lipid probe in a single component bilayer. The rate of protein diffusion is not only lipid dependent but also protein dependent [53], and there is no good framework, of which we are aware, to predict the rate of diffusion of a particular protein in a mixture of lipids. So, a large lipid screen would need to be conducted to optimize the rate of diffusion of KcsA in the membrane. Even, then, it is not clear to what extent this would improve resolution and relaxation.

### 4.2 Effect of Magic Angle Spinning on Samples

Many protocols commonly used during MAS studies, including strong RF irradiation for spectral decoupling and excessive *g*-forces from high spin rates, may be detrimental to proteoliposomes. Biophysical studies of membrane proteins operate on the assumption that the lipid membrane of the model systems is well-defined and does not change over the course of an experiment. The availability and distribution of conformational states may conceivable be affected by these forces. The patency of the lipid bilayer environment is also likely to be influences by these experimental conditions. Organic solids such as proteoliposomes frequently have dense networks of protons, where neighboring proton pair typically have couplings more than 30 kHz. Therefore, experiments often apply MAS rates and RF fields to achieve 100 kHz or more of decoupling to obtain narrow lines.

The centrifugal forces on the sample during MAS described by:

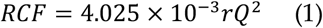

where RCF is given in *g*-force (g), *r* is the radius from the center of the rotor in millimeters and *Q* the spinning frequency in Hz, wherein samples at the rotor wall experience the greatest forces. For example, 3.2 mm rotors typically have a maximum safe spinning frequency of 24 kHz and an inner diameter of 2.17 mm, meaning at top frequency objects at the rotor wall experience a relative centrifugal force 5.0•10^6^ × *g*. The most advanced equipment can result in substantially stronger forces. Even relatively slower spinning frequencies are known to damage biological tissue. Lipid laden mouse adipocytes experience nearly 20% lysis after 2h of MAS at 23 × *g* [56]. Spinning at 20 × *g* for 1 h substantially alters human prostate tissue morphology [47]. As they are denser than their surrounding buffer, proteoliposomes migrate to the rotor wall during MAS, effectively pelleting.

Isopycnic sample preparation by sucrose gradient ultracentrifugation led to highly dilute proteoliposome samples in a high concentration of sucrose which contributes significant background signal to spectra. These are obviously suboptimal conditions for NMR. While not useful for spectroscopy, these experiments did uncover a bimodal distribution of proteoliposome densities. This finding underscores a major theme throughout these studies: the typical way proteoliposome are being prepared for NMR leads to samples are of heterogenous nature, and solid-state NMR samples deserve more scrutiny by complimentary biophysical techniques.

The alternative method of adding highly concentrated sucrose buffer to create a slightly denser then the proteoliposomes to form a “floating pellet” lead to a scenario in which the liposomes migrate to the region where the least amount of centrifugal force is present, namely the center of the rotor, where the *g*-force approaches zero. This allowed for much higher sample concentrations, though still significantly less sample than a fully packed rotor, and with significant small molecule background signal. The lack of dramatic changes to spectra does not justify the use of this technique routinely. However, the slight perturbations we observe suggest that this topic may warrant further investigation.

These studies reiterate previously reported finding from our group that the structurcture of proteoliposome samples is heterogeneous and poorly defined. Future investigations into the effect of magic angle spinning on sample structure would likely benefit from alterative lipid selection. In particular, we have shown that proteoliposomes with DOPC (phosphocholine) headgroups form more predictably spherical structures that would be a better candidate for future investigations [19].

### 4.3 Lipid degradation of KcsA-Proteoliposomes

Work in model liposomes supports previous findings [57] that fatty acid esters can be readily hydrolyzed. In this case, both head groups in the DOPE-DOPS lipid system reacted so slowly that they appeared to be unreacted after two months of incubation on the bench top. It was surprising to find in proteoliposome samples with this lipid composition, evidence for both hydrolysis of fatty acid chains and for lipid headgroups. In one sample, the lipid head groups appear to be entirely hydrolyzed. The catalytic agent leading to hydrolysis in proteoliposome samples has not yet been identified.

Free glycerol ^1^H NMR peaks for the hydrolyzed samples are of the same magnitude as the peaks for free ethanolamine, suggesting that the glycerol peaks do not arise from the co-purifying lipids but rather from the lipids added for reconstitution of liposomes. The most likely cause of the intense glycerol peaks in the ^1^H spectra is that the lipid glycerol backbone is completely hydrolyzed from the phosphate and the fatty acid moieties. Many samples do not show any evidence for free glycerol, but have signal consistent with phosphoglycerol lipid. From this we can conclude that ^13^C-enriched PG co-purifies intact with KcsA in most cases we examined.

This study suggests that proteopliposome samples demand routine quality validation measurements, particularly for studies of the effect of lipid composition. HR-MAS requires specialized equipment and the data acquisition time for an ^1^H-^13^C HSQC is at least 3 hours, making this possible but somewhat onerous. However, here we show that a 1D proton spectrum is adequate to diagnosis the presence of hydrolyzed head groups, which requires no (or very little) change to a CP-MAS experimental setup and can be collected in a matter of minutes.

## 5. Conclusion and Perspective

For many membrane proteins, structure and dynamics information on functionally crucial dynamic loops and termini is lacking. This study demonstrated the use of hybrid solution solid state NMR methods to identify signals for mobile segments of an intrinsic membrane protein in proteoliposomes under functionally relevant conditions. We documented changes in mobility in the C-terminus upon pH triggered activation. We also show evidence that under typical experimental conditions, lipids in proteoliposomes degrade into small molecules with sharp resonances, and demonstrate trivial NMR experiments that can detect the presence of degradation products.

## 6. Methods

### 6.1. Protein Expression and Purification

The fully protonated KcsA data presented in this chapter is from protein that was uniformly ^13^C and ^15^N enriched by expressing KcsA in *Escherichia coli* JM83 cells. The JM83 cell line is a proline auxotrophic, so natural abundance proline was added to cultures.

### 6.2. KcsA Reconstitution

Liposomes were formed from a fixed ratio of lipid by mass with a 9:1 ratio of 1,2-dioleoyl-sn-glycero-3-phosphoethanolamine (DOPE) to 1,2-dioleoyl-sn-glycero-3-phospho-L-serine (DOPS)), which were obtained as a chloroform solution (Avanti), dried as a thin film under N_2_ gas, resolubilized in n-hexane (Sigma), dried again under N_2_ gas and solubilized by bath sonication in 10 mM DM, 50 mM Tris, 100 mM KCl, pH 7.5. Lipids were mixed in mass ratio with KcsA of 1:1, diluted to 2 mM DM and dialyzed in 30 kDa MWCO tubing (Spectrum Chemical) with three exchanges of 4 L of buffer at 12–18 h intervals at room temperature. Proteoliposomes were harvested by centrifugation at 5700 RCF for 30 min and then stored at −80 °C. The presence of KcsA as a tetramer in the liposomes was verified by SDS-PAGE. Pellets were stored at −80 °C until ready for further experimentation.

### 6.3. Lipids

The motivation to use the mixture of lipids in these studies, 9:1 DOPE-DOPS, arose from the previous success of characterizing functional states of KcsA in this particular mixture ^2,23,26^. The PE moiety is zwitterionic and the PS moiety carries a net −1 charge under the conditions used in these pages. Numerous studies have established that lipids with anionic head groups are required for channel activation [58–60], with the presence of those lipids required specifically on the inner leaflet of the membrane [6]. This justifies the use of PS head group in a PE background. Yet there is an entire lipid space that has not been explored in terms of increasing mobility for *J*-based experiments

### 6.4. NMR

*J*-coupled based experiments were performed on a Bruker magnet with a proton field of 750 MHz using a 4 mm high-resolution magic angle spinning probe (HR-MAS) with ^1^H/^13^C/^15^N/^2^H channels with a 40 G/cm gradient coil oriented along the magic angle. Experiments were generally performed between 4–5 kHz MAS and 308 K. Typical hard-pulses were 31kHz for ^1^H, 33 kHz for ^13^C, and 20kHz for ^15^N. Decoupling fields and spinlocks were typically 10 kHz. Heteronuclear decoupling was accomplished using WALTZ16 [61]. HSQCs were phase sensitive using double inept transfer, trim pulses (100 μs), with Echo/Antiecho-TPPI gradient selection and decoupling during acquisition using the Bruker sequence hsqcetgpsi for ^13^C resolved spectra, and hsqcetf3gpsi2 ^57–59^ for ^15^N resolved spectra [62–64]. hCCH-TOCSY spectra were collected with full-rotor period synchronized TOCSY spinlocks, using the Bruker sequence hcchdigp3d2. Site specific *T*_2_ measurements were collected by adding a single rotor-synchronized ^13^C spin-echo between the two inept transfers and increasing the delay over at least five steps until magnetization had decayed to at least 90 %.

Quantitative ^1^H spectra were collected with calibrated ^1^H 90° (typically ~9us), with recycle delays of at least 5s (longest *T*_1_ in samples was typically ~0.9s) with pre-saturation on the H_2_O resonance during the recycle delay with a field strength of approximately 25 Hz with 4 to 16 scans accumulated.

Cross polarization MAS was performed on either the Bruker 750 or a Bruker 900 on 3.2 mm ^1^H/^13^C/^15^N e-free probes at 17 kHz and 19 kHz MAS, respectively and 275 K. Typical field strengths were 100 kHz for ^1^H and 50 kHz for ^13^C. Acquisition times in the direct dimension were approximately 20 ms collected in 2048 points and indirect dimension were typically 4 ms in 128 points. All ^13^C-acquired data was zero-filled to twice the number of points acquired and were multiplied by Lorentzian-to-Gaussian function with 10– 40 Hz of line broadening and 0.3–0.01 Gaussian factors applied empirically in the direct dimension and indirect dimension data were multiplied by sin^2^ function of pure cosine phase.

Solution NMR was performed on a Bruker 500 Ascend instrument using a ^1^H/^13^C/^15^N probe. Sample temperatures were 300 K. Typical field strengths were 18 kHz for ^1^H and 8 kHz for ^13^C for hard pulses, and 2 kHz for ^13^C heteronuclear decoupling using WALTZ16. Homo-soil gradients were accomplished with 7.6 T m^−1^ of 1 ms duration. HSQCs were phase sensitive and multiplicity edited using double INEPT transfer, trim pulses (1 ms), used shaped pulses for inversion on ^13^C (500 μs) with Echo/Antiecho-TPPI gradient selection and decoupling during acquisition (Bruker sequence: hsqcedetgpsp.3). Direct dimensions were acquired for 50 ms in 2048 points and indirect dimensions were acquired for approximately 10 ms in 512 points. The 3D TOCSY data was acquired for 7.5 ms in 128 points in the ^1^H dimension and the ^13^C dimension acquired for 2.5 ms in 64 points. All proton acquired data was zero-filled twice the number of points, rounding up to the nearest perfect-square. FIDs and were multiplied by sin^2^ function of pure cosine phase.

### 6.5. KcsA Cleavage Preparation

To cleave the C-terminus, 1 mg mL^−1^ KcsA in 5 mM decyl-β-maltopyranoside (Anatrace) detergent (DM) was incubated with 20 μg mL^−1^ of bovine α-chymotrypsin (Sigma) for 3 h at 35 °C. KcsA was isolated using His-Select nickel-affinity gel (Fisher), washing with five volumes of buffer 35 mM imidazole and eluting with two volumes of buffer containing 300 mM imidazole. An aliquot of the full-length construct and the post-reaction purified KcsA were analyzed by SDS-PAGE using a 4–12 % Bis-Tris mini gel (Thermo Fisher) at 200 V for 35 min. BLUeye protein ladder (Sigma) was used as a standard. The gel was then stained with PageBlue (Thermo Fisher) coomassie brilliant blue stain according to manufacturer direction. Individual bands were then cut from the gel and placed into new centrifuge tubes and delivered within two hours, on ice, to the proteomics core for mass spectrometry.

### 6.6. Sucrose Gradients

The gradients were prepared by layering equal volumes of two sucrose concentrations (usually 5% and 60 %) in 50 mM Tris, 50 mM KCl, pH 7.25 buffer in 12 mL ultracentrifuge tubes (Beckman), then applying a preprogramed algorithm to mix the layers using a Gradient Master (BioComp Instruments), and cooled at 4 “C overnight. The linearity of the gradient formation protocol was verified by measuring the density of tube fractions using a pipette and balance. Proteoliposome were prepared as described above except a rhodamine-conjugated lipid (1,2-dioleoyl-sn-glycero-3-phosphoethanolamine-N-(lissamine rhodamine B sulfonyl) (Rhod-PE) (from Avanti) was added to the lipid mixture to more easily visualize and quantify the mixtures. The lipid to protein ratio was 1:1 by mass. The lipid mixture was 900:10:1 PE-PS-Rhod PE by mass. 20 mg of KcsA proteoliposomes were suspended in 0.5 mL 5 % sucrose solution then added to the top of the gradient. The tube was subjected to ultracentrifugation, with proper counterweighting, in a Beckman Coulter L70 centrifuge using a swinging bucket rotor (SW41) at 25000 RPM, corresponding to a relative centrifugal field ranging from 47200*g* to 107000*g*, for 24h at 4 °C with the slowest acceleration and no brake during deceleration. After centrifugation, a faint, cloudy band is visible in the lower third of the tube indicating the presence of the proteoliposomes at that layer, this is particularly apparent when Rhod-PE is present, adding a pink tinge to the layer. The bottom of the tube was pierced with a 20 Ga needle that was then removed and the tube was allowed to flow under gravity at 4° C. Fractions of 0.5 mL were collected until the proteoliposome band approached the bottom of the tube when individual drops (~0.1 mL) were collected. Fractions were measured by UV-VIS at 280 nm for the presence of protein and 560 nm (when Rhod PE was used) for the presence of lipids as a qualitative measure. Background scattering caused by the lipids renders 280 nm absorbance sufficient to detect the presence of protein but not to quantify it. To quantify protein in aliquots, we adapted a procedure using bromophenol blue and Triton-X100 from Greenberg and Craddock [9]. Specifically, we formed the assay reagent by mixing 25 mg of bromophenol blue (Sigma), 20 mL of ethanol (HPLC grade, Sigma), 3.0 mL glacial acetic acid (Fischer), 5 mL Triton-X100 (Sigma), and 250 mL deionized water. To implement the assay we mixed the reagent with the analyte in a 9:1 ratio, bath sonicated at 35°C for 5 minutes and measured the UV-VIS absorbance at 610 nm as compared with the regent and deionized water in a 9:1 ratio. We used this reagent to develop a standard curve with KcsA in 1.0 mg/mL in 10 mM DM quantified using UV-VIS at 280 nm [65]. To form the standard curve, we measured 15 concentrations of KcsA solution in triplicate ranging from 0.10 μg mL^−1^ to 20 μg mL^−1^, finding a linear response of the assay to KcsA in this range. We investigated the ability of sucrose to interfere with assay and found no significant difference at sucrose concentrations less than 50 % (w/v). SDS-PAGE), as described above, with silver stain (Pierce silver stain Kit) was used to visualize gradient fraction and verify KcsA remained as a tetramer based on a band appear at approximately 72 kDa as compared with a standard protein ladder (Pierce). The density of each faction of the gradient experiment was measured using an analytical balance of 250 μL of fractions using a micropipette using low retention tips. For highly viscous samples, mass was determined by difference with the solution in the pipette tip.

