## Supplemental information for "High-resolution magic angle spinning NMR of KcsA in liposomes: the highly mobile C-terminus"

**Supplementary Information**

**Sharing these materials is acceptable under CC-BY-NC-ND-4.0.**

**SI Figure 1: SDS PAGE of KcsA C-terminus cleavage**

Full-length (left), following reaction with chymotrypsin (center), and protein ladder (right). Full length tetramer = 70.8 kDa, expected cleavage product = 53.5 kDa.

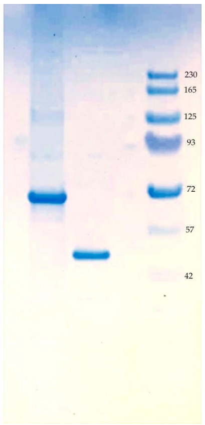


**SI Figure 2: HSQC of KcsA proteoliposomes**

Superimposed $^1H$–$^{13}C$ HSQC by HR-MAS of: full-length KcsA in DOPE-DOPS liposomes (red), KcsA–Δ125 in DOPE- DOPS liposomes (navy), and DOPE-DOPS liposomes (cyan), showing that most resonances in full-length KcsA originate from residues 125-160. Slice of data at dash line ($^{13}C$: 42 ppm) displayed at top, showing PE- ($^1H$: 3.25 ppm) signal intensity is similar for all three samples. All samples: pH 7.25, 50 mM $K^+$ 308 K, 5 kHz MAS.

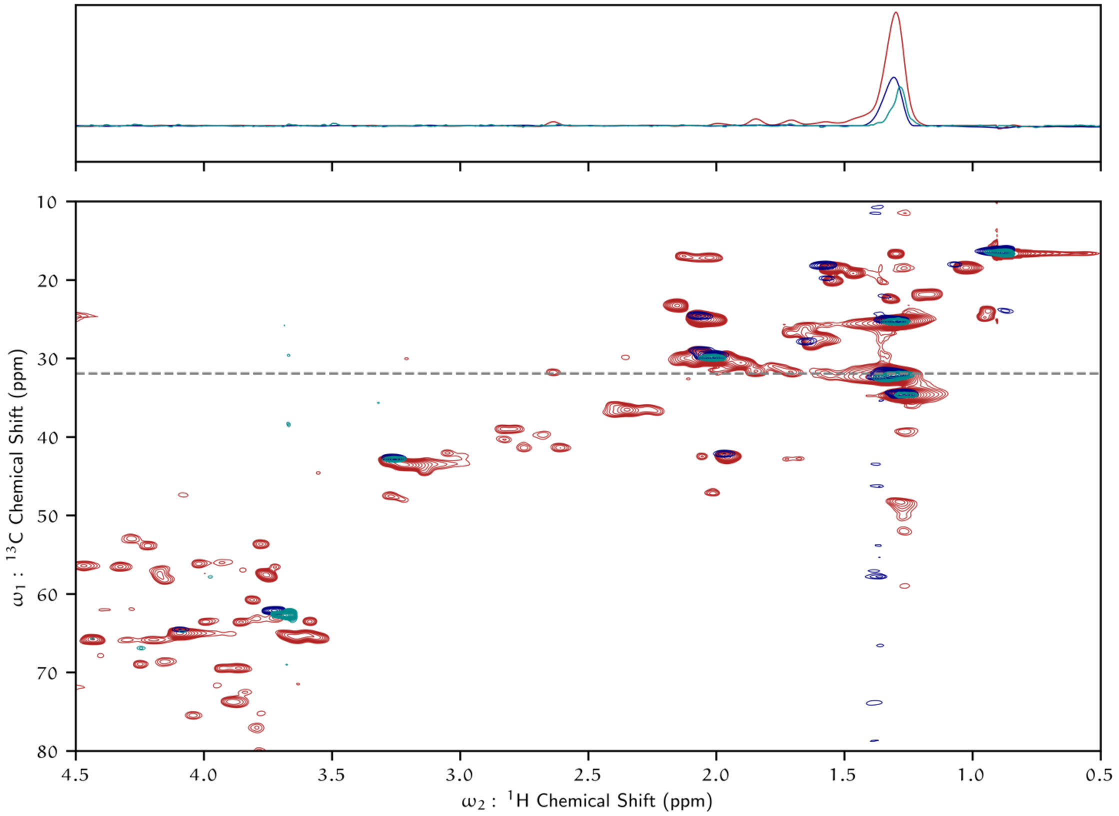

**SI Figure 3: CP-MAS NMR of KcsA proteoliposomes**

Superimposed $^{13}C$–$^{13}C$ proton-driven spin diffusion (DARR) CP-MAS full-length WT KcsA in DOPE-DOPS liposomes (red), KcsA-Δ125 in DOPE-DOPS liposomes (navy), showing the transmembrane domain of KcsA-Δ125 is folded and $^{13}C$ enriched. Assignments displayed are based on prior studies of full-length constructs [23,61] at similar conditions. Slice of data at dashed line ($^{13}C$: 78 ppm) displayed on top. 50 ms DARR mixing period, pH 7.25, 50 mM $K^+$, 270 K, 16.6 kHz MAS.

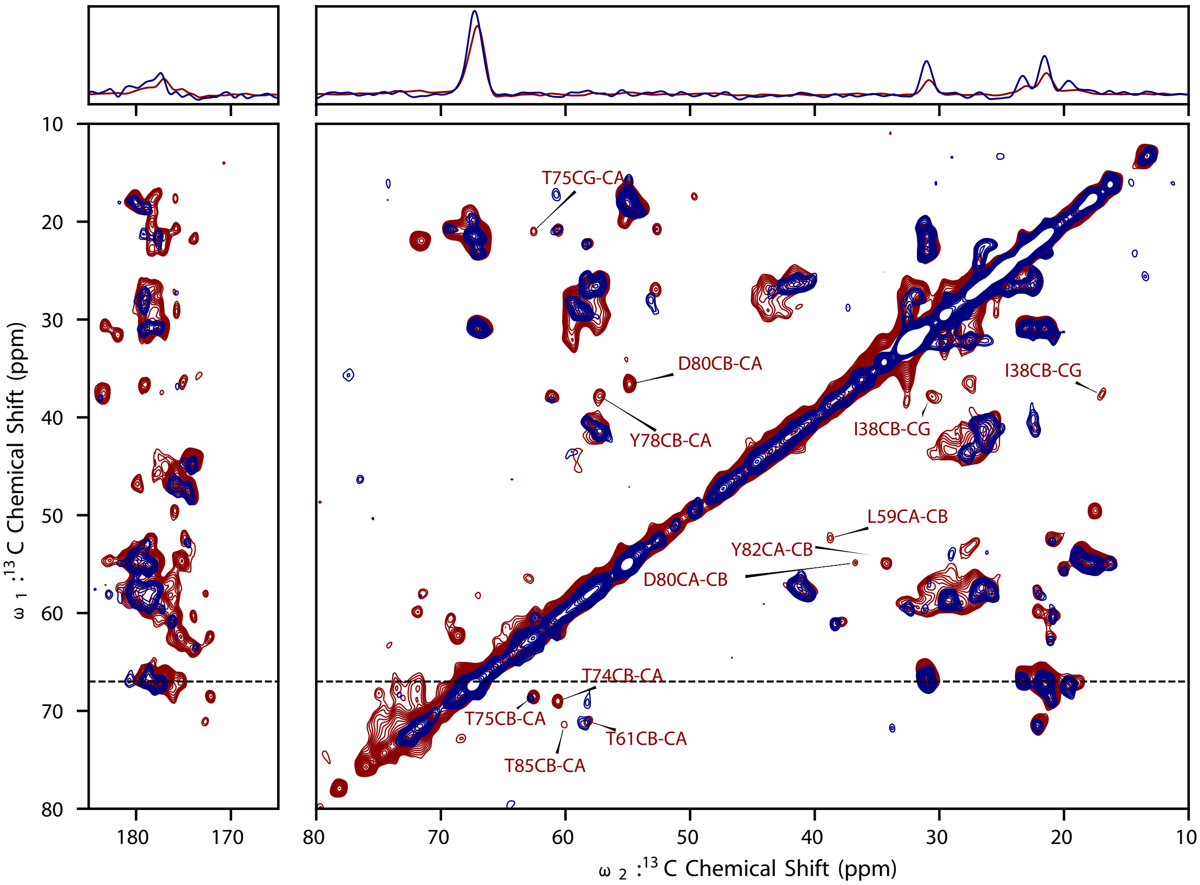


**SI Figure 4: Example of KcsA type assignment from hCCH-TOCSY data**

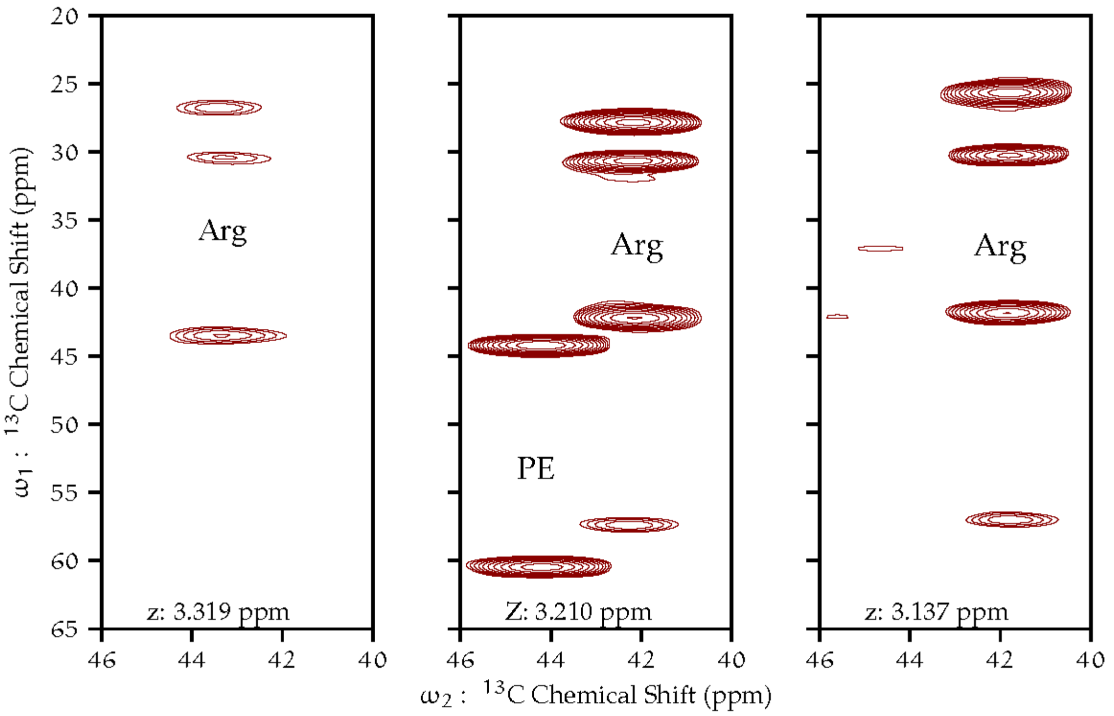


**SI Figure 5: Isopycnic gradient ultracentrifugation of KcsA proteoliposomes**

Silver stained SDS-PAGE gels of aliquots, from two experiments, KcsA ULVs on a sucrose gradient column subjected to ultracentrifugation at 107 000 g (max), with sucrose content indicated.

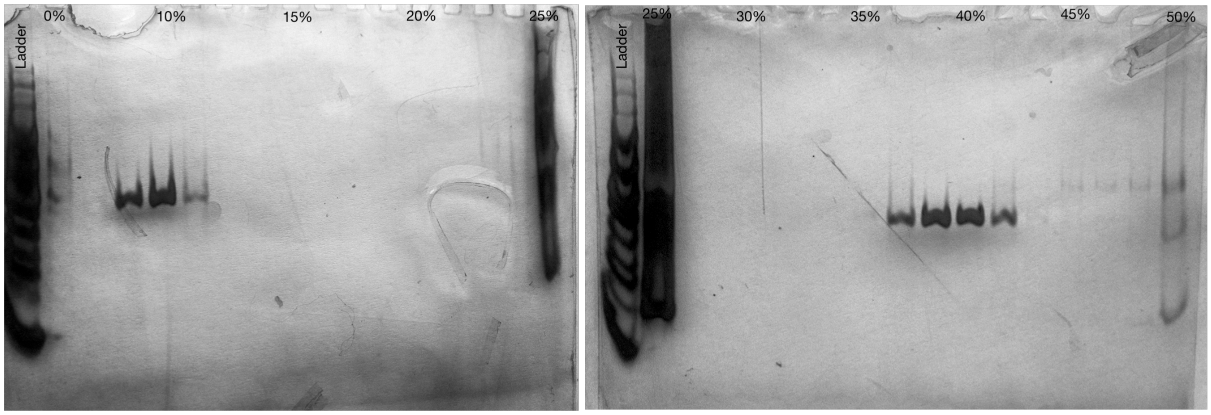


**SI Figure 6: Electron micrographs of KcsA liposomes**

Cryo-electron micrographs of KcsA liposomes. LPR = 1, 9:1 DOPE-DOPS. (A) After 25X freeze thaw (B) Sample 'A' after approximately 160 hours 5kHz MAS 308K, C) Sample 'A' after addition of sucrose, (D) Sample 'C' after approximately 160 hours 5kHz MAS 308K

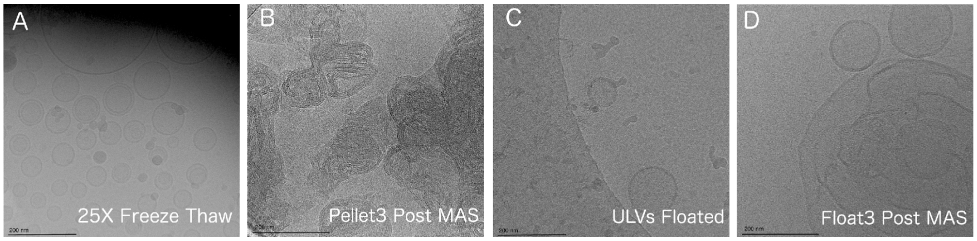


**SI Figure 7: Lipid degredation in KcsA proteoliposome samples from TOCSY Data**

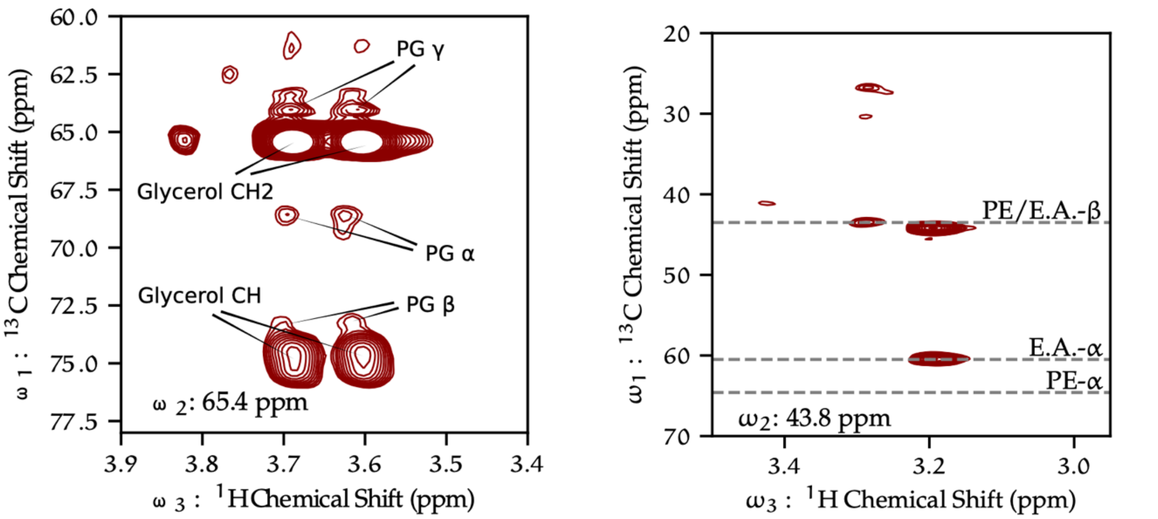


(A) Slice of hCCH-TOCSY 3D correlation at 65.4 ppm. With minor resonances from PG and major from free glycerol labelled.

(B) Slice of hCCH-TOCSY 3D correlation at 43.8 ppm. Expected $^{13}C$ shifts for phospho-ethanol amine related species indicated.

**SI Figure 8: Planes of inter-residue 3D HNCA of KcsA proteoliposomes**

U-13C,15N- KcsA pH 4.0, 50 mM K+. Z-axis (13C (ppm)) range is annotated on each plane. 9 kHz MAS, 308 K.

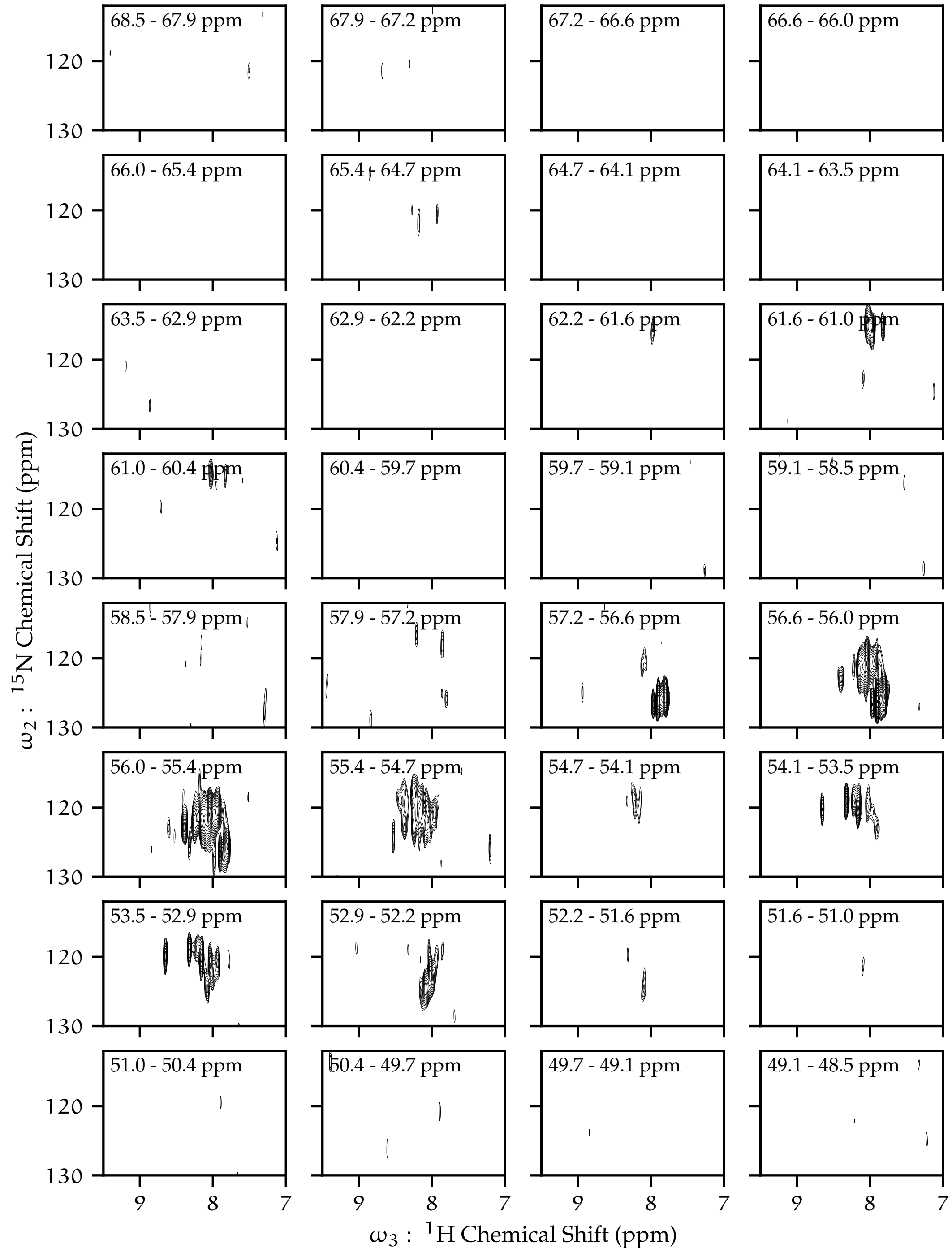


**SI Figure 9: Planes of intra-residue (i+1) 3D HNCO of KcsA proteoliposomes**

Fractionally deuterated U-13C,15N- KcsA at pH 4.0, 50 mM K+. Z-axis (13C (ppm)) range is annotated on each plane. 9 kHz MAS, 308 K

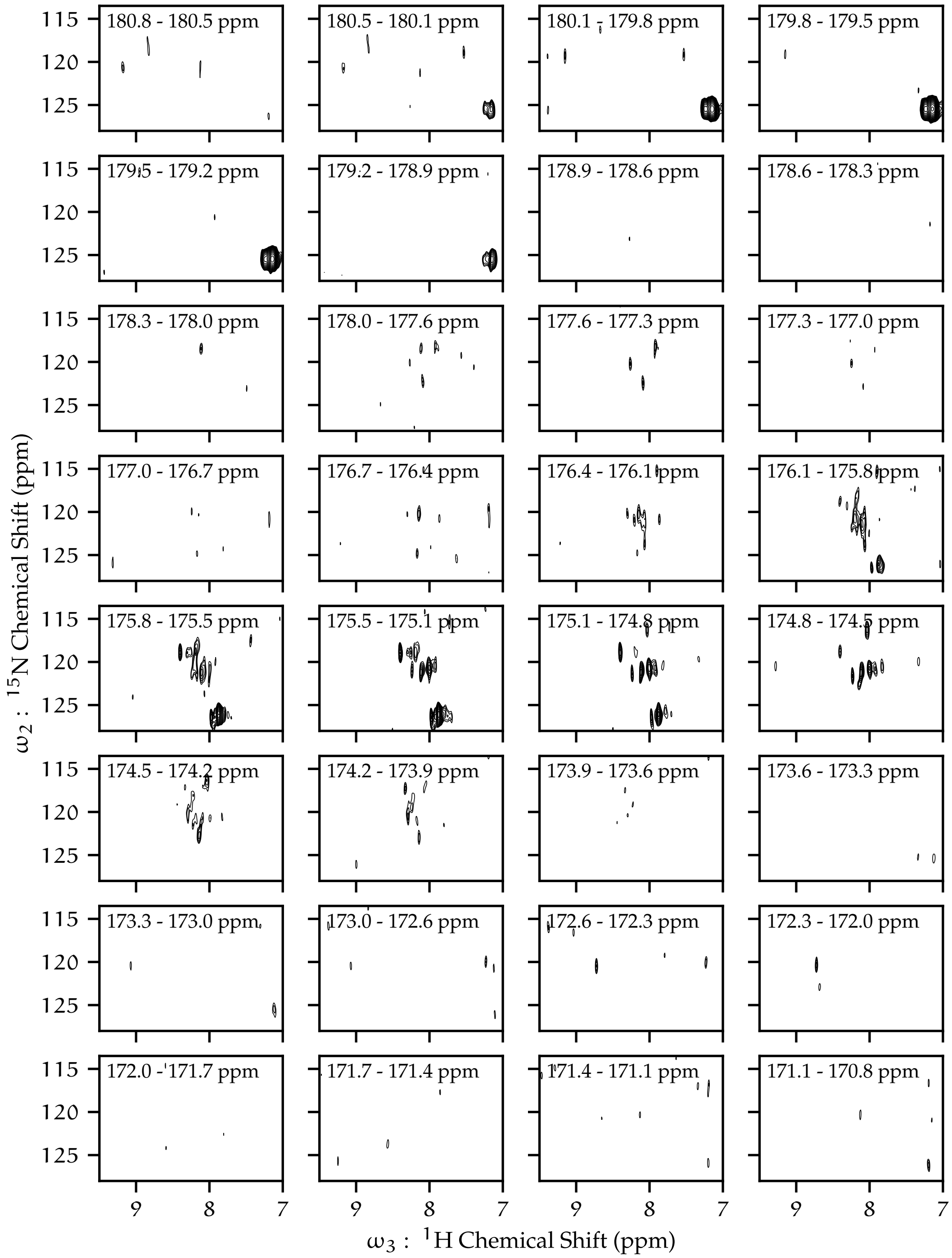

180.8 - 180.5 ppm
180.5 - 180.1 ppm
180.1 - 179.8 ppm
179.8 - 179.5 ppm
179.5 - 179.2 ppm
179.2 - 178.9 ppm
178.9 - 178.6 ppm
178.6 - 178.3 ppm
178.3 - 178.0 ppm
178.0 - 177.6 ppm
177.6 - 177.3 ppm
177.3 - 177.0 ppm
177.0 - 176.7 ppm
176.7 - 176.4 ppm
176.4 - 176.1 ppm
176.1 - 175.8 ppm
175.8 - 175.5 ppm
175.5 - 175.1 ppm
175.1 - 174.8 ppm
174.8 - 174.5 ppm
174.5 - 174.2 ppm
174.2 - 173.9 ppm
173.9 - 173.6 ppm
173.6 - 173.3 ppm
173.3 - 173.0 ppm
173.0 - 172.6 ppm
172.6 - 172.3 ppm
172.3 - 172.0 ppm
172.0 - 171.7 ppm
171.7 - 171.4 ppm
171.4 - 171.1 ppm
171.1 - 170.8 ppm
115
120
125
9
8
7
$\omega_2$ : $^{15}$N Chemical Shift (ppm)
$\omega_3$ : $^{1}$H Chemical Shift (ppm)

### SI Figure 10: Residue types identified in hCCH-TOCSY data of KcsA (Table 1) mapped on to KcsA sequence

KCSA with His6 tag

N-TERMINUS

| 1 | | | | | | | 2 | 3 | 4 | 5 | 6 | 7 | 8 | 9 | 10 | 11 | 12 | 13 | 14 | 15 | 16 | 17 | 18 | 19 | 20 |
|---|---|---|---|---|---|---|---|---|---|---|---|---|---|---|---|---|---|---|---|---|---|---|---|---|---|
| M | H | H | H | H | H | H | P | P | M | L | S | G | L | L | A | R | L | V | K | L | L | L | G | R | H |

TRANSMEMBRANE DOMAIN

| 21 | 22 | 23 | 24 | 25 | 26 | 27 | 28 | 29 | 30 | 31 | 32 | 33 | 34 | 35 | 36 | 37 | 38 | 39 | 40 | 41 | 42 | 43 | 44 | 45 | 46 | 47 | 48 | 49 | 50 | 51 | 52 | 53 | 54 | 55 | 56 | 57 | 58 | 59 | 60 | 61 | 62 | 63 | 64 | 65 | 66 | 67 |
|---|---|---|---|---|---|---|---|---|---|---|---|---|---|---|---|---|---|---|---|---|---|---|---|---|---|---|---|---|---|---|---|---|---|---|---|---|---|---|---|---|---|---|---|---|---|---|
| G | S | A | L | H | W | R | A | A | G | A | A | T | V | L | L | V | I | V | L | L | A | G | S | Y | L | A | V | L | A | E | R | G | A | P | G | A | Q | L | I | T | Y | P | R | A | L | W |

| 68 | 69 | 70 | 71 | 72 | 73 | 74 | 75 | 76 | 77 | 78 | 79 | 80 | 81 | 82 | 83 | 84 | 85 | 86 | 87 | 88 | 89 | 90 | 91 | 92 | 93 | 94 | 95 | 96 | 97 | 98 | 99 | 100 | 101 | 102 | 103 | 104 | 105 | 106 | 107 | 108 | 109 | 110 | 111 | 112 | 113 | 114 | 115 |
|---|---|---|---|---|---|---|---|---|---|---|---|---|---|---|---|---|---|---|---|---|---|---|---|---|---|---|---|---|---|---|---|---|---|---|---|---|---|---|---|---|---|---|---|---|---|---|---|
| W | S | V | E | T | A | T | T | V | G | Y | G | D | L | Y | P | V | T | L | W | G | R | L | V | A | V | V | V | M | V | A | G | I | T | S | F | G | L | V | T | A | A | L | A | T | W | F | V |

C-TERMINUS

| 116 | 117 | 118 | 119 | 120 | 121 | 122 | 123 | 124 | 125 | 126 | 127 | 128 | 129 | 130 | 131 | 132 | 133 | 134 | 135 | 136 | 137 | 138 | 139 | 140 | 141 | 142 | 143 | 144 | 145 | 146 | 147 | 148 | 149 | 150 | 151 | 152 | 153 | 154 | 155 | 156 | 157 | 158 | 159 | 160 |
|---|---|---|---|---|---|---|---|---|---|---|---|---|---|---|---|---|---|---|---|---|---|---|---|---|---|---|---|---|---|---|---|---|---|---|---|---|---|---|---|---|---|---|---|---|
| G | R | E | Q | E | R | R | G | H | F | V | R | H | S | E | K | A | A | E | E | A | Y | T | R | T | T | R | A | L | H | E | R | F | D | R | L | E | R | M | L | D | D | N | R | R |

Residues identified in hCCH-TOCSY data

| D/N | F | K | L | R |
|---|---|---|---|---|
| 2 | 2 | 1 | 2 | 3 |

**SI Table 1: KcsA $^{13}C$ $T_2$s**

$^{13}C$ α $T_2$ U-$^1H$,$^{13}C$,$^{15}N$-KcsA in 9:1 DOPE-DOPS liposomes, pH 4.0, 50 mM $K^+$, 308K, 5 kHz MAS. Lipids assigned from chemical shifts, other resonances assumed to be protein H-Cα resonances

| Site | Type | $^1H$ (ppm) | $^{13}C$ (ppm) | $^{13}C$ $T_2$ (ms) |
|---|---|---|---|---|
| | protein | 4.04 | 51.4 | 3.6 |
| | protein | 3.98 | 54.6 | 3.4 |
| Arg-α | protein | 4.03 | 55.2 | 2.8 |
| Met-α | protein | 4.30 | 55.8 | 3.9 |
| | protein | 4.16 | 56.9 | 2.7 |
| Lys-α | protein | 3.72 | 56.9 | 2.8 |
| | protein | 4.37 | 61.6 | 2.2 |
| Thrβ | protein | 4.19 | 69.0 | 2.4 |
| PE-β | lipid | 3.24 | 43.9 | 6.5 |
| PE-α | lipid | 4.05 | 64.2 | 5.9 |
| PG-γ | lipid | 3.56 | 64.7 | 3.7 |
| PG-γ | lipid | 3.63 | 64.7 | 3.8 |
| PG-α | lipid | 3.82 | 68.9 | 4.3 |
| PG-α | lipid | 3.88 | 68.9 | 4.4 |
| PG-β | lipid | 3.80 | 73.1 | 7.2 |

### SI Table 2: $^{1}$H-$^{13}$C HSQC per sample chemical shifts and relative intensities of various species

$^{1}$H-$^{13}$C HSQC chemical shifts of exogenous lipid signals in KcsA liposome samples by HR-MAS. "–" indicates no detected signal for given peak and "o" indicates the presence of grossly overlapping peaks in peak region. : glycerol $CH_2$ is degenerate with PG γ signal. All samples reconstituted in natural abundance phospholipids with dioleoyl fatty acids chains with lipid-to-protein ratio of 1 (w/w), 308 K, 5 kHz MAS, internally referenced to DSS. Sample details given in SI Table 3.

| ID | Protein | Lipid | pH | PG α<br>$^{1}$H (ppm) / $^{13}$C (ppm) | PG β | PG γ & Glycerol $CH_2$† | Glycerol CH | HC=C | PE-β (%) | EA-β (%) | PG-β (%) | gly-β (%) |
|---|---|---|---|---|---|---|---|---|---|---|---|---|
| A | (empty) | PE/PS | 7.3 | – | – | – | – | – | 100 | 0 | 0 | 0 |
| B | KcsA | PE/PS | 7.3 | 3.94/73.7 ; 3.88/68.4 | 3.94/68.5 | 3.69/65.4 ; 3.61/65.4 | 3.83/74.9 | 5.1/126.9 | 42 | 57 | 21 | 79 |
| D | KcsA | PE/PS | 6.3 | 3.92/69.4 ; 3.86/69.4 | 3.89/73.7 | 3.63/65.4 ; 3.61/65.3 | – | 5.09/127.1 | 100 | 0 | 100 | 0 |
| E | KcsA | PE/PS | 4 | 3.92/69.6 ; 3.85 / 69.6 | 3.89/73.8 | 3.66/65.2 ; 3.6/65.2 | – | – | 100 | 0 | 100 | 0 |
| F | $^{2}$H KcsA | PE/PS | 6.3 | – | – | 3.64/65.5 ; 3.55/65.5 | – | – | 100 | 0 | 100 | 0 |
| G | $^{2}$H KcsA | PE/PS | 4 | – | 3.95/73.6 | 3.71/65.4 | 3.8/74.5 | – | 26 | 74 | 27 | 73 |
| H | KcsA–ΔC | PE/PS | 7.3 | – | – | – | – | – | 100 | 0 | 0 | 0 |
| I | KcsA | PE/PS | 7.3 | 3.94/69.1 ; 3.89/69.1 | 3.93/73.5 | 3.67/65.4 ; 3.58/65.4 | o | – | 100 | 0 | – | – |
| J | KcsA | PE/PS | 7.3 | 3.95/69.4 ; 3.88/69.3 | 3.92/73.6 | o | o | – | 100 | 0 | – | – |
| K | KcsA | PE/PS | 7.3 | o | o | 3.66/65.4 ; 3.57/65.4 | o | – | 85 | 15 | – | – |
| L | KcsA | PE/PS | 6.3 | 3.98/69.3 ; 3.91/69.3 | 3.96/73.6 | 3.7/65.5 ; 3.61/65.4 | o | 5.11/126.9 | 100 | 0 | – | – |
| M | KcsA | PC | 4 | 3.92/73.8 | 3.96/69.5 | 3.7/65.3 ; 3.63/65.3 | – | 5.05/127 | 0 | 0 | 100 | 0 |

**SI Table 3: KcsA proteoliposome sample details**

Description of various samples used in HR-MAS samples, providing details for **SI Table 3:** H-$^{13}$C HSQC chemical shifts

| | Protein | Labeling | Lipids | pH | sucrose (% w/w) | buffer | $K^+$ (mM) | $Mg^{2+}$ (mM) | $D_2O$ (% vol.) |
|---|---|---|---|---|---|---|---|---|---|
| A | no protein | | DOPE/DOPS | 7.3 | 0 | tris | 50 | 0 | 15 |
| B | KcsA | U-$^{13}$C$^{15}$N | DOPE/DOPS | 7.3 | 0 | tris | 50 | 0 | ~90 |
| D | KcsA | U-$^{13}$C$^{15}$N | DOPE/DOPS | 6.3 | 0 | pipes | 50 | 0 | 10 |
| E | KcsA | U-C$^{13}$C$^{15}$N | DOPE/DOPS | 4 | 0 | citrate | 50 | 0 | 10 |
| F | KcsA | F–$^{2}$H U-$^{13}$C$^{15}$N | DOPE/DOPS | 6.3 | 0 | pipes | 50 | 0 | 10 |
| G | KcsA | F–$^{2}$H U-$^{13}$C$^{15}$N | DOPE/DOPS | 4 | 0 | citrate | 50 | 0 | 7 |
| H | KcsA–ΔC | U-$^{13}$C$^{15}$N | DOPE/DOPS | 7.3 | 0 | tris | 50 | 0 | ~90 |
| I | KcsA | U-$^{13}$C$^{15}$N | DOPE/DOPS | 7.3 | 38 | tris | 50 | 2 | 10 |
| J | KcsA | U-$^{13}$C$^{15}$N | DOPE/DOPS | 7.3 | 42 | tris | 50 | 0 | 10 |
| K | KcsA | U-$^{13}$C$^{15}$N | DOPE/DOPS | 7.3 | 44 | tris | 50 | 2 | 10 |
| L | KcsA | U-$^{13}$C$^{15}$N | DOPE/DOPS | 6.3 | 45 | tris | 50 | 0 | 10 |
| M | KcsA | U-$^{13}$C$^{15}$N | DOPC | 4 | 0 | citrate | 150 | 0 | ~20 |
